# Axial Nephron Fate Switching Demonstrates a Plastic System Tunable on Demand

**DOI:** 10.1101/2025.03.29.646044

**Authors:** MaryAnne A. Achieng, Jack Schnell, Connor C. Fausto, Réka L. Csipán, Matthew E. Thornton, Brendan H. Grubbs, Nils O. Lindström

## Abstract

The human nephron is a highly patterned tubular structure. It develops specialized cells that regulate bodily fluid homeostasis, blood pressure, and urine secretion throughout life. Approximately 1 million nephrons form in each kidney during embryonic and fetal development, but how they develop is poorly understood. Here we interrogate axial patterning mechanisms in the human nephron using an iPSC-derived kidney organoid system that generates hundreds of developmentally synchronized nephrons, and we compare it to in vivo human kidney development using single cell and spatial transcriptomic approaches. We show that human nephron patterning is controlled by integrated WNT/BMP/FGF signaling. Imposing a WNT^ON^/BMP^OFF^ state established a distal nephron identity that matures into thick ascending loop of Henle cells by endogenously activating FGF. Simultaneous suppression of FGF signaling switches cells back to a proximal cell-state, a transformation that is in itself dependent on BMP signal transduction. Our system highlights plasticity in axial nephron patterning, delineates the roles of WNT, FGF, and BMP mediated mechanisms controlling nephron patterning, and paves the way for generating nephron cells on demand.

## INTRODUCTION

Each nephron that forms during embryonic and fetal development follows a deeply conserved developmental program that, by birth, ensures fluid bodily homeostasis^1^. To generate a nephron, mesenchymal nephron progenitors (NPs) gradually cluster together into pretubular aggregates (PTAs), that in turn epithelialize into renal vesicles (RVs), later giving rise to the tubular Comma and S-shaped body nephron stages (CSB and SSB)^2,3^. It is during this process that the early nephron generates a proximal-to-distal (PD) axis, a symmetry breaking event that over time produces ∼24 functionally specialized cell-types spatially distributed along the adult nephron PD axis^1,4^. The proximal-most end of the adult nephron consists of podocytes that generate a physical filter interacting with capillaries in the glomerular tuft while the remainder of the nephron is a tubular structure consisting of several broad segments - proximal tubule, descending loop of Henle, ascending Loop of Henle, macula densa, distal convoluted tubule (DCT), and connecting tubule (CNT), each divisible into subtypes and by function^4–7^. At the distal-most end, the nephron forms a luminal connection to drain urine into the collecting duct. While fate-mapping has shown all adult mouse nephron cells form from NPs^8^, the mechanisms underpinning the gradual specification of PD identities are poorly understood, particularly in humans.

Identifying mechanisms underlying human nephron patterning requires *in vitro* systems that replicate the development of the nephron PD axis. Many studies have produced kidney organoids with nephron-like structures (nephroids), generated by directed differentiation from pluripotent stem cells. However, resulting nephroids primarily reflect developmental rather than mature cell states and distal nephron segments are poorly represented^9–11^. To overcome these limitations, it is essential to understand and mimic distal nephron development *in vitro*.

*In vivo*, distal and proximal nephron identities form sequentially as NPs are recruited into the PTA, RV, and CSB^3^. Several signal inputs regulate distal nephron development^12–14^. The distal domain is marked by transcription factors HNF1B and POU3F3^3,15^, and their mouse orthologs regulate development of nascent nephron epithelia and maturation of distal segments of loop of Henle, macula densa and distal tubule^16–18^. Distal cell identities require an integration of dosed Wnt, Bmp, Notch, and Fgf signaling^19–22^ and perturbations to these pathways generate distinct phenotypes. However, interpretations of these phenotypes are hindered by the nephron’s complex morphogenetic nephrogenic program and dynamic gene expression patterns. Wnt9b likely initiates nephrogenesis by driving β-catenin-mediated differentiation and cell aggregation to form PTAs^19,23–27^. Thereafter, distal and proximal positional identities are sensitive to the dosage of β-catenin as increasing or decreasing the transcriptional output of β-catenin distalizes and proximalizes nephrons, respectively^28^. While it is clear that β-catenin is central to nephron PD patterning, our previous work in mice shows that nephron cells can be further sensitized to β-catenin by simultaneously modulating Bmp or Pi3k signaling, with each combination generating distinct axial patterning outcomes^28^.

In this study, using the WNT and BMP signal transduction pathways as a starting point, we develop an approach to generate hundreds of nephroids that reproducibly break their PD axis symmetry and generate defined nephron cell identities. Using a combination of single cell sequencing, small scale organoid screening, spatial transcriptomics, and pathway perturbations, we show that distal identities are dependent on conserved patterning programs in the human iPSC-derived nephron. By examining the expression dynamics of FGF8 *in vivo* during early nephron development, we link the ligand’s expression to the forming ascending loop of Henle and demonstrate these cell-fates depend on FGF signaling. Strikingly, distal fated nephron cells devoid of FGF signaling revert to generate proximal cell identities, a program that we in turn show is conditional on BMP signal transduction. The system as whole demonstrates considerable cell fate plasticity and provides a model for switching human nephron cell fates at will, and defines the spatiotemporal requirements for WNT, FGF, and BMP signaling during axial nephron patterning.

## RESULTS

### Spontaneous nephron axial patterning in a malleable window

To establish a system for controlling human nephron cell identities *in vitro*, we adopted and modified a kidney organoid protocol^29^ and identified a differentiation window for manipulating PD axial symmetry breaking during the PTA to RV transition. Briefly, human iPSCs are differentiated in monolayer culture for 7 days to mimic WNT-mediated posterior paraxial mesoderm induction and FGF-mediated intermediate mesoderm induction, before being transitioned into sheet-like 3D organoid cultures (**Figure 1a**). At differentiation day 10 (DD10), these WT1^+^ organoids display synchronously epithelializing PTA-like structures forming apical-basal polarities (**Figure S1a-c**). An apical surface positive for tight-junction protein ZO-1 faces inwards while the outside of each PTA deposits a basal β-laminin^+^ basement membrane (**Figure 1b, Figure S1b**). At DD10, PTAs are positive for NP marker SIX1 and display uniform distribution of transcription factors and PD patterning markers PAX2, PAX8, LEF1, and Notch ligand JAG1, and are surrounded by MEIS1^+^ interstitium (**Figure S1a-c**). Over a 2-day period, each nephroid thereafter elongates and morphologically begins to resemble RVs, displaying upregulation of JAG1 and epithelial cadherin CDH1 (**Figure 1b**). At this stage, a portion of nephrons break synchronicity and mimic the polarized downregulation of WT1 observed *in vivo*. Compared to DD10 when 482/482 (100%) of nephroids are symmetric along their PD axes, on DD12, 36/368 (10%) nephroids display polarized downregulation of WT1, highlighting the first timepoint a PD axis and asymmetry is observed *in vitro* (**Figure 1b**). In this early mosaic state, the majority (90%) of nephroids maintain uniform levels of WT1 protein, but by DD14, 88/383 (23%) of nephroids have WT1^-^ domains (**Figure 1b, c**). When the PD axis develops *in vivo*, distinct CDH1^+^ distal, JAG1^+^ medial, and WT1^+^ proximal domains form (**Figure 1b**); CDH1 marks precursors of the distal nephron, JAG1 marks proximal/medial nephron cells, and WT1 is found in podocytes and parietal epithelium, respectively. *In vitro*, however, DD14 nephroids only partially recapitulate this marker segregation and the majority of nephrons – 295/383 (77%) – are positive for WT1 along their whole axial polarity. Additionally, while human SSBs establish sharp boundaries between each PD domain, CDH1, JAG1, WT1 boundaries in organoid nephroids overlap. Consistent with the broad detection of WT1 at precursor stages (DD14), nephroids are broadly positive for podocyte marker MAFB and differentiate into large MAFB^+^ renal corpuscles (DD28), surrounded by a CD31^+^ vasculature-like networks; few GATA3^+^ distal tubules form (**Figure S1d-e**). This indicates that the nephroids in this system display a bias towards podocyte-forming programs (**Figure 1d**).

**Figure 1.**
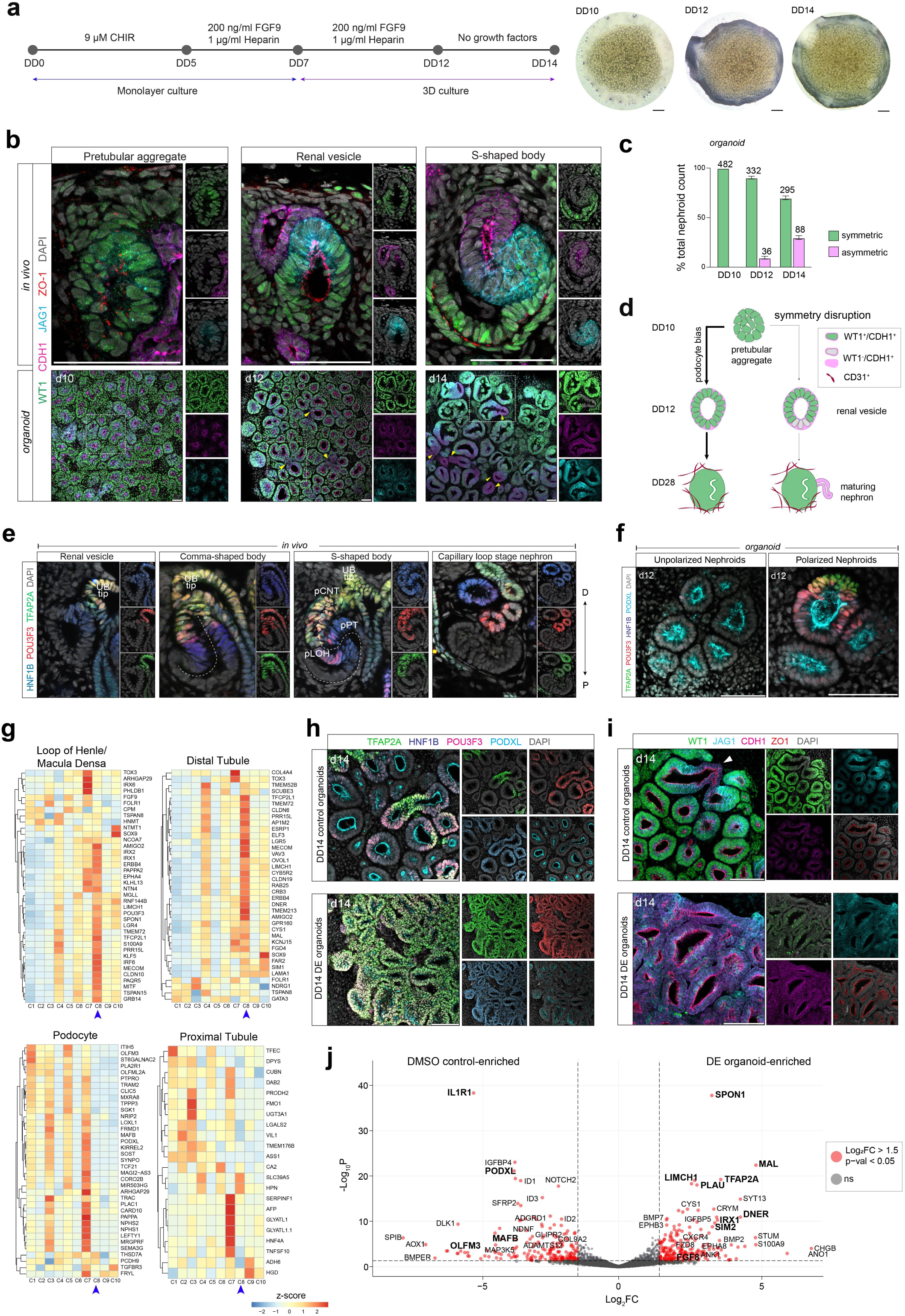
Directed differentiation of kidney organoids for a proximal-to-distal fate switch. (a) (left) Schematic of modified Takasato protocol for iPSC-derived kidney organoids. (right) Brightfield images of DD10, DD12 and DD14 organoids. Scale bar: 500 microns. (b) Human kidney immunofluorescence stains for proximal, medial, and distal domain markers at the PTA, RV and SSB stages. Corresponding stage-matched kidney organoids at DD10, DD12 and DD14 are stained for the same markers. Yellow arrows denote polarized nephrons. Scale bar: 50 microns. (c) Bar graph showing percentage of total nephrons per organoid symmetrical vs asymmetrical WT1 expression on DD10, DD12 and DD14. Numbers above graph denote average nephroids counted per organoid (n = 2). (d) Schematic of kidney organoid differentiation trajectory from DD10-DD28. (e) Immunofluorescence stains of human kidneys at the RV, CSB, SSB and CLSN stages showing transcription factors marking the distal nephron domains. The SSB includes labeled positional relationships of precursors of the CNT (pCNT), DCT (pDT), loop of Henle (pLOH), proximal tubule (pPT) and podocyte (pPod). (f) Immunofluorescence stains of DD12 organoids showing distal transcription factor expression in polarized vs unpolarized nephrons. Scale bar: 50 microns. (g) Heatmaps of TPM value z-scores of nephron cell type markers at the SSB stage, expressed across 10 differentiation conditions in DD14 organoids (n = 2). Blue arrow denotes condition 8. (h-i) Immunofluorescence stains of DD14 control (condition 1) vs DE (condition 8) organoids showing proximal-distal marker expression. Scale bar: 50 microns. (j) Volcano plot of DESeq analysis comparing DD14 DMSO control and DE organoid total mRNA sequencing data. Dashed lines show 0.05 p-value cutoff (horizontal) and ±1.5-fold change cutoff (vertical).

In human nephrogenesis, transcription factors HNF1B, POU3F3 and TFAP2A define distal nephron precursors. HNF1B is detected at the distal-most end of the early renal vesicle and, together with POU3F3 and TFAP2A, is thought to mark early to late distal segments (**Figure 1e**). As the SSB elongates, domains of these transcription factors shift, forming 3 distinct segments from the distal-most to medial SSB: HNF1B^+^/POU3F3^+^/TFAP2A^+^, HNF1B^+^/POU3F3^+^/TFAP2A^-^ and HNF1B^high^/POU3F3^-^/TFAP2A^-^ regions. These segments correspond to putative precursors of the distal nephron, segment 3 of the proximal tubule, and segments 1 and 2 of the proximal tubule, respectively^1^. In contrast, as most nephroids have not broken symmetry at the RV stage (DD12; see **Figure 1c**), few nephroids express these 3 distal transcription factors and rare triple HNF1B^+^/POU3F3^+^/TFAP2A^+^ domains are observed (**Figure 1f**, **Figure S1f**). This is consistent with the infrequent distal nephron phenotype seen at DD28. These views suggest that cell type outcomes are established during the initial stages of differentiation and highlight a time frame (DD10-DD12) for manipulating axial nephron patterning and reading outcomes at DD14.

### Altering WNT and BMP signals shifts nephron axial patterning to distal programs

To generate distal axial nephron cell fates, we constructed a dosage-duration-timing matrix increasing canonical WNT signaling and reducing BMP signaling during the PTA to RV transition. We have previously linked β-catenin signaling to positively regulate distal axial identities in the mouse nephron^28^, while BMP/SMAD reporters are active in proximal axial identities^28,30^.

We activated WNT signaling using GSK3β inhibitor, CHIR99021 (CHIR), a well-studied WNT agonist, modulating dosages and duration and timing dosing to avoid disrupting the mesenchymal-to-epithelial transition (MET)^19,23,31^. We inhibited BMP signaling using LDN 193189 dihydrochloride (LDN), a selective inhibitor of BMP receptors ALK2 and ALK3 that prevents phosphorylation of SMAD1/5/8^32^. To promote NP epithelialization during the PTA-to-RV transition, we incorporating PI3K inhibitor, LY2940021 (LY29), from DD10 to DD12^28,33^. We assayed patterning outcomes at DD14 by whole-mount immunofluorescent stains for PD markers (in PD order: WT1, PODXL (also marks apical surfaces together with ZO1), JAG1, HNF1B, CDH1, POU3F3, TFAP2A) and by bulk RNA sequencing at DD14 and DD20 to determine long term patterning, across these 2- and 3-inhibitor combinations (**Figure S1g**).

In combination, conditions 4, 6, 7, 8, 9, and 10 (**Figure S1g**) displayed evidently larger TFAP2A^+^ domains relative to control (cond. 1), CHIR-only (cond. 2), LDN-only (cond. 3) and LY29-only (cond. 5) treatments (**Figure S1h**). This agrees with bulk RNA sequencing data where on their own, each small molecule inhibitor (SMI) only negligibly affected axial domain shifts (**Figure S1i**). WNT activation from DD11-DD12 yielded a modest increase in distal domain emergence relative to control organoids (slight upregulation of POU3F3, HNF1B and TFAP2A), but this ineffectively suppressed podocyte (WT1^+^) differentiation (**Figure S1i**). This is consistent with our previous *ex vivo* data demonstrating a transient and reversible effect of SMI pulse treatments when used in solitarily^28^. To increase resolution to the precursor populations generated in each SMI condition, we assessed DD14 and DD20 organoid expression of genes (top 50) that are normally enriched in axially distributed PD nephron precursors in *in vivo* SSBs, as described in Lindström et al^1^ (**Figure 1g**, **Figure S1j, k**). While no *in vivo* SSB precursor gene list was fully represented, meeting the cutoff criteria of TPM > 10, podocytes (37/50 genes), loop of Henle (38/50 genes) and distal tubule (37/50 genes) precursors had the highest representation in our organoids, while fewer parietal epithelium-, proximal tubule- and connecting tubule-enriched genes were expressed (31/50, 22/50 and 25/50 genes, respectively). Condition 8 (DD10-11 PI3K inhibition, DD11-12 PI3K inhibition, WNT activation, BMP inhibition), henceforth referred to as distal-enriched (DE), generated nephron-like precursors resembling those of the DCT, loop of Henle, and CNT at the SSB stage (DD14), and maintained expression of genes while simultaneously and suppressing podocyte-enriched genes until DD20 (**Figure 1g**, **Figure S1j, k**). Consistent with these findings, DE organoids expressed the highest level of TFAP2A and POU3F3 (**Figure S1l**). Integration of PI3K inhibition, WNT activation, and BMP inhibition was thus identified as a candidate condition for generating DE nephroids.

Direct comparisons between control and DE nephroids showed a remarkable switch from proximal to distal axial identities. DE nephroids lost their proximal/podocyte (WT1^+^) and gained distal (TFAP2A^+^/POU3F3^+^/HNF1B^+^) cell profiles (**Figure 1h-i**). TFAP2A activation was systemic in DE organoids and was strongly co-detected with POU3F3 and HNF1B (**Figure 1h**). While WT1 expression was prominent in control organoids, it was suppressed in DE organoids, which instead elongated forming CDH1^+^/JAG1^low^ nephroid tubules (**Figure 1i**). An equivalent direct comparison of the total mRNA signature of DD14 control and DE organoids (**Table S1**) further illustrates this dramatic shift in cell identities. Control organoids express podocyte-enriched genes such as *OLFM3, PODXL, MAFB,* and *IL1R1*, while DE organoids are enriched for distal nephron markers, *TFAP2A, MAL, IRX1, SIM2, BMP7* and *SPON1* (**Figure 1j**)^1,34^. Together, these data provide strong evidence that simultaneous activation of WNT/β-catenin and suppression of BMP/SMAD signal transduction synergistically drives nephroid distalization.

### A transient BMP^OFF^ and WNT^ON^ state primes nephroid cells for distal nephron formation

To assess the axial shift of our DE nephroid model, we performed single cell RNA sequencing to analyze the cellular composition of DD14 DE and control organoids. As expected, both control (8219 cells) and DE (9301 cells) organoids consisted of nephron-like (*PAX2*^+^) or interstitial-like (*PDGFRA^+^*) cells, with few off-target cell types (no detection of muscle marker, *MYOD1*) (**Figure S2a, b; Table S2**). Both nephron and interstitial progenitors are derived from the intermediate mesoderm^35–37^, indicating our organoid cell composition efficiently generates intermediate mesoderm and nephron cells. Control organoids contained a small subset of CDH5^+^ endothelial progenitors (**Figure S2a**), which likely give rise to the CD31^+^ vasculature-like cells observed in DD28. Nephron-like cells were the dominant cell-population in the organoids with 80% (6650 cells) and 77% (7158 cells) of cells from control and DE organoids, respectively, being classified as nephron lineage (**Figure 2a**).

**Figure 2.**
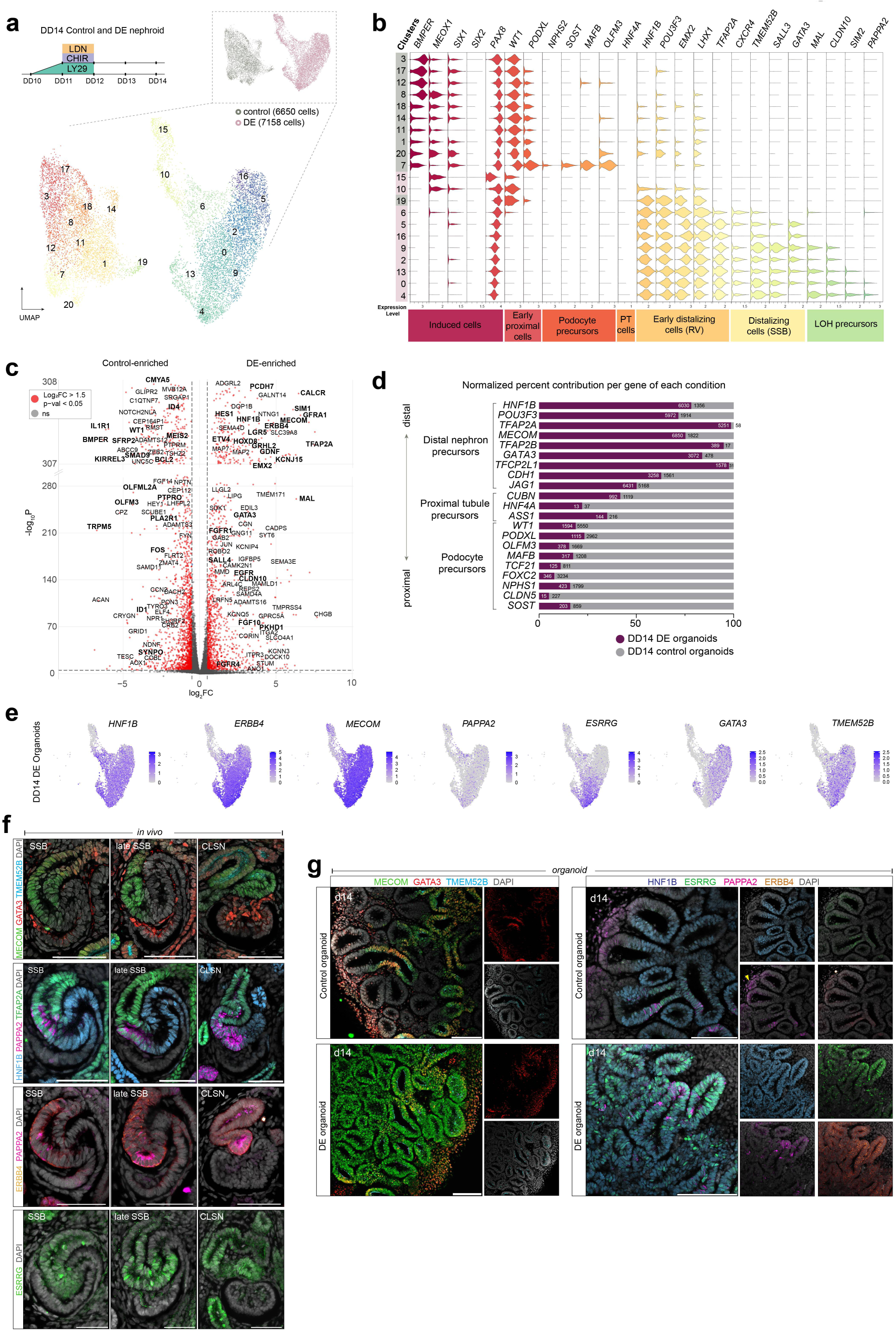
Single-cell RNA sequencing data shows nephron lineage divergence between control and DE organoids. (a) Annotated unsupervised clustering of PAX2^+^ nephron-like DD14 control and DE organoid cells represented in a UMAP. (Inset) Sample contribution of each condition. (b) Violin plot with annotations of select gene markers (top x axis) showing expression levels (bottom x axis) in the DD14 organoid dataset. The y axis shows cluster numbers and sample origin (gray square – control organoids, pink square – DE organoids). (c) Volcano plot of DESeq analysis comparing DD14 DMSO control and DE organoid nephron scRNA sequencing data. Dashed lines show 0.05 p-value cutoff (horizontal) and ±1.5-fold change cutoff (vertical). (d) Bar plot showing normalized percent contribution of cells expressing select nephron markers (log_2_FC > 0) separated by sample origin. Numbers in bars show raw cell counts. (e) Feature plots of select genes in the DD14 DE organoid dataset. (f) Immunofluorescence stain of week 16 human kidney from SSB-CLSN stage marking DCT/CNT precursor and loop of Henle/macula densa precursor segments. (g) Immunofluorescence stains of DD14 control and DE organoids showing expression of DCT/CNT precursor and loop of Henle/macula densa precursor markers. Yellow arrow – autofluorescence. Scale bar: 50 microns.

When control and DE nephroid cells were merged and compared, few cells co-clustered with the other condition, suggesting minimal overlap in equivalent cell states between the 2 conditions (**Figure 2a**, top right). Cell states clustered based on differentiation states and cellular processes defined by nephron patterning markers (**Figure 2a, b; Table S2**). Clusters 3, 8, 12 and 17, comprised of control nephroid cells, displayed expression of induced nephron progenitor markers, *BMPER* and *LHX1,* while *PAX8,* another induction marker, was expressed across both conditions. As expected, control cells broadly expressed *WT1* across all clusters while DE nephroid cells were largely devoid of *WT1* transcripts, expressed only in clusters 10 and 15 (**Figure 2b, Figure S2c**). Controls also displayed remnant expression of NP markers, *SIX1* and *MEOX1*. *Bona fide* differentiation of podocytes (*NPHS2^+^, PODXL^+^, SOST^+^, TCF21^+^*) was only observed in controls and the *WT1*^+^ DE cells did not express these markers, indicating podocyte differentiation was suppressed, alluding to rudimentary remnants of proximal-like cells. Instead, DE nephroid cells were strongly biased towards distal axial identities expressing *TFAP2A*^+^ (**Figure S2d**), consistent with our immunofluorescence data. These *TFAP2A^+^* cells co-expressed a range of early distal RV markers like *EMX2, LHX1,* and *POU3F3* and late distal RV markers, *GATA3, MAL* and *CXCR4*^1,3,6,7,15^. A portion of DE cells also expressed differentiation markers (*CLDN10, SIM2* and *PAPPA2*) normally enriched in the loop of Henle and macula densa of the *in vivo* nephron^6,7^ (clusters 13, 0 and 4; **Figure 2b**).

To scrutinize differences in gene expression of control and DE nephroid cells we performed comparative differential gene expression analyses, identifying 480 control and 629 DE enriched genes (p<0.05, log2FC>1.5; **Figure 2c**; **Table S1**). These data corroborate conclusions that control organoids generate a podocyte-like gene signature positive for *WT1*, *IL1R1*, *KIRREL3, OLFM3, PTPRO, PLA2R1, SYNPO,* while DE nephroids adopt a broad and distinct distal-biased profile exhibiting expression of markers indicating early distal nephron development (*LGR5*, *CALCR*, *MAL, TFAP2A, HNF1B, MECOM, GATA3*) and precursor specification and maturation (*TMEM52B, ERBB4, ESRR*γ, *PAPPA2*) (**Figure 2c, S2e; Table S1**). The overt shift in differentiation towards distal nephron identities was further emphasized by quantifying cells expressing PD axial markers. DE nephroid cells were strongly biased to develop distal nephron identities while control nephroid cells generate proximal-like cells (**Figure 2d**). At DD14, neither condition contained many *HNF4A*^+^ cells, but control nephroids had ∼4x more *HNF4A*^+^ cells than DE organoids.

Control nephroids displayed enrichment of BMP/SMAD pathway effectors *ID1*, *ID2*, *ID3*, and *ID4*^38^ (**Figure 2c; Figure S3a; Table S1**) compared to DE nephroid equivalents. To evaluate BMP activity, we assayed active SMAD signaling by immunolabelling dual phospho-SMAD1/5. Strong phospho-SMAD labelling was detected in control nephroids and in their surrounding interstitium, but this was absent in the DE organoids, consistent with LDN efficiently suppressing BMP/SMAD signaling in DE conditions (**Figure S3b**). While the role of BMP/SMAD signaling is not established in the human nephron, *in vivo* labelling of developing nephrons indicated phospho-SMAD activity in proximal PTAs, RVs, CSB, and in the proximal precursors of the SSB and CLSN (**Figure S3c**), thus further correlating BMP signaling with human proximal axial nephron identities as suggested in mice^39^. Conversely, DE organoids showed upregulation of canonical WNT targets *JUN*, *DLL1, CDH1, EMX2, SALL4, TFCP2L1* and *WNT4*^40–43^ (**Figure S3d**), as expected from the addition of CHIR increasing WNT/β-catenin signaling.

DE nephroids expressed genes indicating specification of maturing distal axial identities. Within *HNF1B*^+^, *TFAP2A*^+^, *ERBB4*^+^, *MECOM*^+^ DE nephroids cells, select cells were also positive for *PAPPA2*, *ESRRG*, *GATA3*, and *TMEM52B* (**Figure 2e, Figure S2e**). In developing human nephrons, MECOM marks the entire distal nephron domain in the SSB, GATA3 and TMEM52B mark distal precursors of the CNT and DCT at the distal-most parts of the nephron, while the maturing ascending loop of Henle and macula densa precursors are marked by PAPPA2, ESRRγ and ERBB4; HNF1B and TFAP2A are included for reference (**Figure 2f**). ESRRγ, a receptor that acts as a transcriptional activator in the absence of a bound ligand, marks loop of Henle/macula densa precursors^44^ and is detected in their nuclei in the SSB in a mosaic pattern. The receptor ERBB4 is first detected at the SSB stage in the distal limb. Its detection coincides with that of metalloprotease, PAPPA2, a putative marker of loop of Henle and macula densa precursors. Using these as anchors to scrutinize the DE nephroid differentiation, immunolabelling demonstrated that DE nephroids are shifted to a distal MECOM^+^/TMEM52B^+^ cell state that is broadly co-labeled with loop of Henle receptor ERBB4, and mosaic detection of maturation transcription factor ESRRγ and macula densa/loop of Henle marker, PAPPA2 (**Figure 2g**). These markers are rare in control nephroids and suggest that the DE nephroids are driven towards a distal tubule fate with a further sub-specification towards the distal tubule, macula densa, and thick ascending limb of the loop of Henle. Interestingly, both organoid conditions display sporadic GATA3^+^ domains. This specification agrees with unbiased GO term analyses of genes upregulated in DD14 DE organoids (log_2_FC > 1.5) (**Figure S3e, Table S1**); DE genes include those associated with nephron epithelium development, calcium ion binding, and ion transmembrane transporter activity, consistent with functions associated with the distal nephron epithelia. Of note, FGF receptor binding was highlighted as a predicted enriched molecular function in DE organoids, attributed to significant upregulation of *FGF8*, *FGF10* and *FGF18*, indicating elevated FGF signaling in DE organoids relative to controls.

To ensure the DE protocol is robustly reproducible, we performed experiments replacing either the PI3K inhibitor LY294002 with GDC-0941, a structurally different and potent PI3K inhibitor, or replaced SMAD inhibitor LDN-193189 with Dorsomorphin, a BMP inhibitor that targets BMP receptors, ALK2, ALK3 and ALK6^45–47^. Both conditions replicated results of the DE protocol, displaying a global shift to distal axial identities marked by elongated distal TFAP2A^+^, CDH1^+^ tubules with apical PODXL^+^ surfaces, positive for ERBB4, and increased abundance of PAPPA2^+^ cells (**Figure S4a**). To further benchmark the DE protocol, we generated and distalized kidney organoids from a different iPSC HNF4A-YFP reporter line^48^. These iPSCs distalized and displayed near-global expansion of HNF1B, POU3F3 and TFAP2A axial identities with co-activation of β-catenin target LEF1, and mosaic expression of GATA3, while downregulating podocyte (WT1, MAFB), and proximal tubule precursor markers (HNF4A^+^) (**Figure S4b-e**). In contrast, controls displayed a bias towards forming WT1^+^/MAFB^+^/ZO1^high^ podocyte-like cells and few HNF1B^+^/POU3F3^+^/TFAP2A^+^ domains. Our validation experiments show that the DE protocol is adaptable to different cell lines and SMIs.

These observations support the hypothesis that driving distal cell identities (WNT/β-catenin activation) and suppressing proximal cell differentiating (BMP/SMAD inhibition) generates distal nephron-like cells. Moreover, these data present a distal nephron-like organoid model enriched with loop of Henle and DCT precursor-like cells.

### Distalized nephrons differentiate to loop of Henle fates

To directly identify the closest *in vivo* equivalents of DE nephroid cell states, we generated an *in vivo* transcriptome framework by reanalyzing previous single cell RNA sequencing capturing human nephrogenesis data from week 14 and week 17 human kidneys ^1,49^, processed as described in Schnell et al^33^. In these data, NPs (*MEOX1^+^/CITED1^+^*) develop into podocyte (*MAFB^+^*), parietal epithelium (*PLAT^+^*), proximal tubule (*HNF4A^+^/SLC3A1^+^*), and early distal tubule precursors (*HNF1B*^+^/*POU3F3*^+^/*TFAP2A*^+^) that differentiate into loop of Henle (*SLC12A1^+^/PAPPA2^+^/CLDN10^+^*) and a combined cluster consisting of DCT and CNT cells (*TMEM52B^+^/GATA3^+^/EMX1^+^*) (**Figure 3a**, **Fig S5a-b**, **Table S2**). Using these as a reference, we mapped control and DE nephroid transcriptomes to the most similar *in vivo* cell RNA profiles (**Figure 3a**). DE nephroids closely resembled distal precursor cells, with a minor WT1^+^ NP-like cell population remaining in these organoids. Strikingly, DE nephroid cells also mapped to loop of Henle precursors, but not DCT and CNT precursors (**Figure 3a**, **S5c**). Control nephroid cells closely corresponded to PTA and early podocyte cells, and to a lesser degree, the earliest PT precursors, where proximal and distal cell fates have not fully diverged (**Figure 3a, S5c**). Cell type prediction scores between nephroid and *in vivo* counterparts showed that DE organoids most closely resemble the distal nephron precursor state and loop of Henle precursors, while control nephroids most resemble the PTA-like state and early podocytes (**Figure 3b**). There is therefore a strong prediction that DE nephroid cells are in an early distal nephron identity programmed towards a loop of Henle-like state.

**Figure 3.**
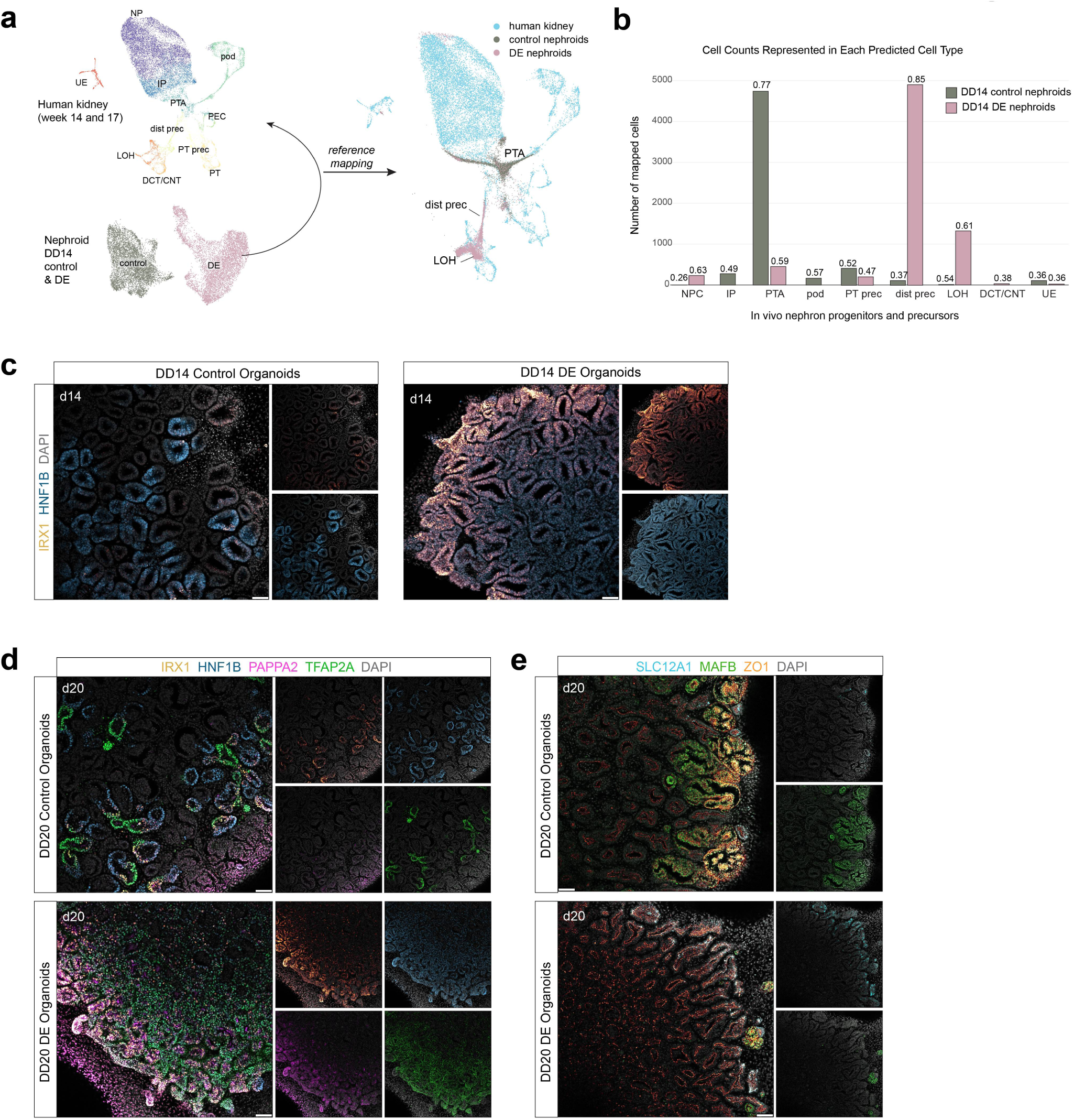
Characterizations of distal nephron segmentation in distal-enriched organoids. **A.** schematic showing reference mapping of DD14 organoid scRNA sequencing dataset to reanalyzed week 14 and week 17 merged human nephron and ureteric scRNA sequencing data ^1,49^. (Right) UMAP of human kidney (reference) overlaid with control and DE organoid mapping. **B.** Bar plot showing number of organoid cells mapping to each predicted cell type, split by condition. Numbers above bars denote prediction score for each cell type. (c) Immunofluorescence stain of DD14 control and DE organoids showing detection of IRX1. (d-e) Immunofluorescent stains of DD20 control and DE organoids showing detection of select loop of Henle/macula densa precursor and podocyte precursor markers. Abbreviations: NP – nephron progenitors, IP – induced progenitors, PTA – pretubular aggregate, pod - podocytes, PEC – parietal epithelial cells, dist prec – distal nephron precursors, PT prec – proximal tubule precursors, PT – proximal tubule, LOH – loop of Henle, DCT/CNT – distal convoluted tubule/connecting tubule.

To increase resolution to potentially primed differentiation programs in the DE model and their transcriptional drivers, we benchmarked expression of all known human transcription factors in *in vivo* nephrons (approach described in Kim, S., et al^6^). We identified 39 distal nephron-enriched transcription factors that have either very low or no expression in other nephron cell types (see Methods) (**Figure S5d**); *IRX1*, *IRX2*, *TFAP2B*, *SIM2, IRX6, TFEB* and *NRK* are enriched in loop of Henle/macula densa precursors, *EMX1*, *GATA3*, *PRDM16* and *NR0B2* are enriched in distal convoluted tubule and connecting tubule precursors, and *GATA2, ELF5* and *EHF* are enriched in the ureteric epithelium (UE). Several of these transcription factors are known to be distal nephron-enriched, and co-expression of *Irx1*, *Irx2* and *Sim2* in loop of Henle/macula densa precursors is conserved in P0 mouse nephrons^6^, indicating function. Analysis of transcription factors in our DE nephroids showed that loop of Henle-enriched transcription factors, *IRX1*, *IRX2*, *TFAP2B* and *SIM2,* are restricted to *PAPPA2*^+^ DE nephroid clusters 13, 0 and 4 (**Figure S5e**). Of the DCT-enriched transcription factors, only *GATA3* was detected. *EMX1*, which is normally expressed in the DCT and CNT precursors of the CLSN, was absent. Transcription factors that are predicted to be expressed in more mature loop of Henle segments, *IRX6, TFEB* and *NRK,* and DCT segments, *NR0B2, PRDM16* and *EMX1*, were not detected. Their lack of detection in DD14 nephroids, which resemble the SSB stage, was expected. We observed strong co-expression of known loop of Henle/macula densa precursor-enriched markers, *IRX1* and *SIM2,* in a subset of DE nephroid cells (**Figure S5f**). Validation experiments show IRX1 is broadly present in DD14 DE nephroids (**Figure 3c**), indicating these cells are initially driven to a distal axial state primed to thereafter generate a subset of more mature nephron cell types.

Our bulk RNA sequencing of DD14 and DD20 DE organoids show that the gene signatures of loop of Henle and macula densa lineages are maintained and reinforced, indicated by large increases in expression of *PAPPA2*, *ERBB4,* and ascending loop of Henle/macula densa solute carrier, *SLC12A1* (**Figure S5g**), while transcription factors *IRX1* and *SIM2* expression were maintained (**Figure S5g**). Validation by immunostaining in DD20 organoids shows strong detection of IRX1, PAPPA2, and SLC12A1 in DE organoids (**Figure 3d-e**). In contrast, maturation markers of the DCT such as *EMX1,* and calcium transport-associated *TRPV5* and *CALB1* were undetected in both DD14 and DD20 DE organoids, while *GATA3* expression remained the same between the two time-points and *SALL3* expression was downregulated (**Figure S5g**). Podocyte signatures did not recover by DD20, where DE organoids displayed few MAFB^+^ cells, relative to controls (**Figure 3e**).

We further assessed differentiation outcomes at DD28. DE organoids consisted primarily of TFAP2A^+^/SLC12A1^+^/ERBB4^+^/CDH1^+^ cells that were mosaically positive for PAPPA2 and IRX1 (**Figure S5h, S6a**). Markers of other nephron axial identities, e.g., proximal tubule (HNF4A^+^), podocyte (MAFB+), and CNT (GATA3+) were rare (**Figure S6b**). In contrast, control organoids were enriched with MAFB^+^ glomeruli-like cells abutting CD31^+^ vasculature and RENIN^+^/GATA3^+^ juxtaglomerular-like cells, contained HNF1B^+^/HNF4A^+^ proximal tubule-like cells, and displayed sparse PAPPA2 and IRX1 labeling (**Figure S6c**); RENIN^+^ cells were not observed in DE organoids. Controls also displayed sustained activation of genes associated with mature podocyte markers, *ITIH5* and *THSD7A*^44,49,50^, glomerular endothelial cell marker *PLVAP* and VEGF receptor *FLT1* (**Figure S6d**, **Table S1**). DE organoids further upregulated genes encoding distal-nephron-specific receptors and Na^+^ channels, *CLDN16*, *KCNJ1*, *CASR*, *SLC12A1* and *SCNN1A,* and known distal nephron markers, *MAL*, *CDH1* and *TFAP2A*, However, as has been observed with long-term cultures of kidney organoids^10^, off-target genes emerged, such as muscle-associated *MSC*, and *MYOZ2*, and neuron markers, *NEUROD1* and *NPTX2*. GO term analyses of top differentially expressed genes in whole DD28 DE organoids underline distal nephron-linked functions of ionic exchange (**Figure S6e**). Together, these data demonstrate a model for a nearly systemic axial switch in nephron cell type identities.

### Modulation of FGF and BMP signaling shifts domain identities along the proximal-distal nephron axis

The DE organoid approach demonstrates that simultaneous activation of WNT and inhibition of BMP/SMAD signaling generate cells primed for distal nephron fates. DE organoids significantly increased *FGFR1, MAP7* and *MAP2* expression at DD14 (**Figure 2b**, **Table S1**), highlighting a potential role of FGF/MAPK signaling in distalization, in line with mouse nephron studies showing *Fgf8* is required for distal nephron cell survival and differentiation and is a direct target of Wnt/β-catenin^23,51^.

Consistent with this view, *FGF8* was strongly upregulated in DE nephroids (**Figure 4a**; log_2_FC ∼3), as was *FGF10* (**Figure 2c**, **Figure S7a**). *In vivo*, *FGF8* is upregulated during distal nephron development in a sequence closely matching *TFAP2A* expression, but *FGF8* is turned off soon after SSB stages (**Figure 4b**). *FGF9* is upregulated later in distalization when *FGF8* is downregulated, and *FGF10* follows a similar onset of expression to *FGF8* but persists in the DCT and CNT precursors (**Figure 4b; Figure S7b**). To resolve the exact sequence of *FGF8* up and downregulation, we generated spatial transcriptomic data capturing human kidney and nephron development^52^. *FGF8* is detected in *TFAP2A*^+^ distal cells in the early RV, remains there into the late RV, and then expands proximally within the *TFAP2A^+^*domain (**Figure 4c**). By SSB stages, *FGF8* is detected in the forming *IRX1^+^* domain and is downregulated in distal-most *GATA3^+^* cells (**Figure S7c**). By late SSB stages, *FGF8* is downregulated throughout the distal *TFAP2A^+^* cells aside from the *IRX1^+^* and *PAPPA2^+^* cells, and a subset of *FGF8^+^* cells express loop of Henle marker *SLC12A1*. *FGF8* transcripts are not detected in *HNF4A*^+^ proximal and *MAFB*^+^ podocyte precursors (**Figure S7c**).

**Figure 4.**
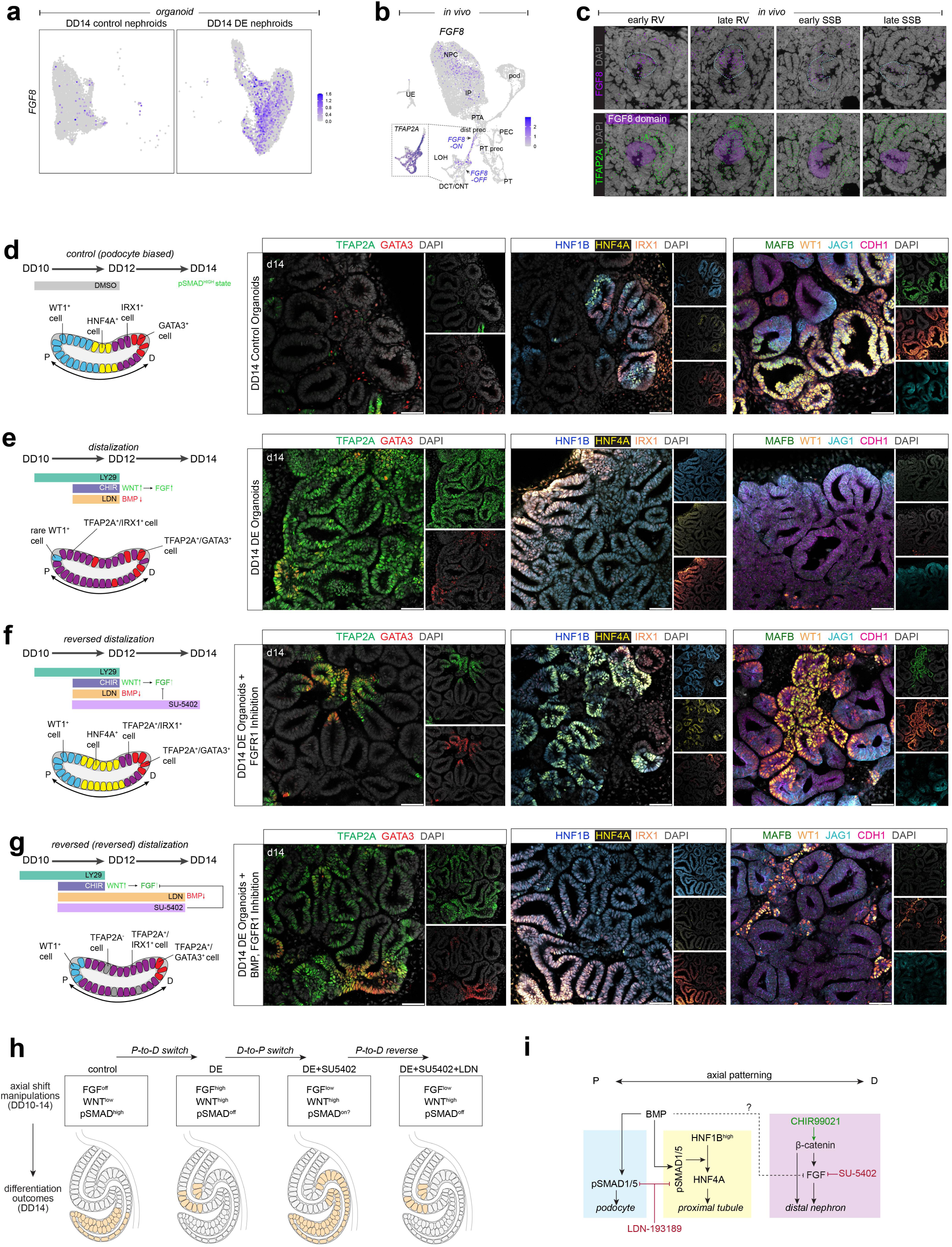
Dynamics of FGF, WNT and BMP signaling in nephron segmentation. (a) Feature plot of FGF8 expression in DD14 organoids, split by condition. (b) Feature plot of FGF8 in the human kidney dataset. Inset: TFAP2A detection in the distal nephron. (c) SeqFISH data of week 16 human kidney showing detection of FGF8 and TFAP2A from early RV-late SSB. (d-g) (left) Schematics of SMI treatment and nephron segmentation in DD14 organoids and (right) immunofluorescent stains showing detection of DCT/CNT, proximal tubule, loop of Henle/macula densa and podocyte precursor markers (right). Scale bars: 50 microns. (h) Schematic of nephron segments and the corresponding pathways that drive their development. (i) Proposed schematic of select signaling pathway interactions in the developing mammalian nephron.

Considering potentially paired receptors for FGF8, Fgfr1 is detected in the nascent nephron and tubular epithelia of mice^51^. Our single cell analyses of the human nephron show *FGFR1* is enriched in NPs, induced NPs, PTAs, and proximal and distal axial lineages, but is at lower levels in podocytes (**Figure S7d**). *FGFR2* is enriched in the parietal and ureteric epithelia, while the onset of *FGFR3* is highest in the maturing proximal tubule (**Figure S7d**). In DD14 organoids, *FGFR1* expression is detected in both control and DE organoids, though at higher levels in the latter (**Figure S7e**, **Figure 2c**, **Table S1**). Low levels of *FGFR2* are also detected in all clusters of both conditions, except for DE organoid cluster 4, that is enriched with loop of Henle precursor markers (**Figure S7e**, **Figure 2b**). Interestingly, mirroring *in vivo* nephrons, *FGFR3* is only detected in control organoid cluster 19 (**Figure S7e**), the only cluster with HNF4A^+^ cells (not shown). As the preferred receptor for FGF8, FGFR1 is therefore a candidate receptor for FGF signaling in the distal nephron.

To determine whether FGF signaling driven by β-catenin activation is responsible for distal nephron development, we inhibited FGFR1 between early RV- to SSB-like stages (DD11-DD14) using SU-5402, a potent FGFR1 inhibitor^53,54^, and compared forming nephroids with control and DE conditions (**Figure 4d-f**). This condition generated a Wnt/β-catenin active state unable to signal via FGFR1. Strikingly, inhibition of FGFR1 in DE organoids reversed the DE phenotype (**Figure 4f**). Distal nephron marker TFAP2A was almost entirely blocked, IRX1 reduced, and conversely, HNF4A^+^ proximal tubule domains yet again formed next to MAFB^+^/WT1^+^ podocytes (**Figure 4f**; compared to d and e). Moreover, HNF4A^+^ cells formed within HNF1B^high^ domains, strongly reminiscent of normal *in vivo* development where elevated HNF1B expression in the medial SSB drives HNF4A^17,33,55^.

These data show that inhibition of FGFR1 in DE organoids results in a next-to-complete reversal of the DE state back to a proximal axis-biased differentiation outcome. Since HNF4A^+^ proximal tubule precursors and MAFB^+^ podocytes formed in the DE+FGFR1-inhibitor condition, we therefore asked whether this switch is dependent on SMAD signaling driving proximal axial cell-fates again, when FGF8 is rendered incapable of distalizing nephroids due to FGFR1 inhibition. To test this, we extended the SMAD inhibitor treatment with LDN so that DE organoids simultaneously received the FGFR1 inhibitor and the SMAD inhibitor until DD14 (**Figure 4g**). Remarkably, this again flipped the differentiation outcome so that TFAP2A and IRX1 were detected throughout nephroids, WT1 and MAFB were lost, and HNF1B was again detected at a homogenous level (as in DE conditions) without HNF4A. While TFAP2A was detected in a mosaic pattern, IRX1^+^ cells were detected throughout nephroids. Of note, we observed only a modest expansion of the HNF4A domain in control organoids treated with SU-5402 for the same duration as the other experiments, despite expressing *FGFR1*, (**Figure S7e-f**) indicating that it is the DE condition that generates a FGF8^high^ state driving a distal fate. When analyzing differentiation endpoints (DD28) of DE and DE+FGFR1 inhibitor treated organoids, the latter developed podocytes as well as loop of Henle identities, showing that the axial shifts imposed by DD14 generate nephroids that, thereafter, differentiate towards their respective primed states (**Figure S7g-h**).

Altogether, these data demonstrated that the nephron PD polarity is a highly tunable epithelial axis, and molecular functions required for nephron cell type specification can be disentangled in organoid models, thus assembling recipes for differentiating cell-fates along the PD axis (**Figure 4h**). Our model proposes pathway-mediated differentiation to form the different nephron segments and maintain boundaries of their domains (**Figure 4i**). WT1 cooperatively functions with MAFB in activation of mature podocyte transcription factors and thus plays a critical role in podocyte formation^56–58^. WT1^+^ podocyte domains expand in the presence of Tankyrase inhibitor, IWR1^28^, suggesting a requirement for suppressed β-catenin activity for podocyte formation. From our and others’ exploration of FGF signaling in nephrons, FGFR1/Fgfr1-mediated signaling drives distal nephron formation. Evidence of FGF and WNT cooperative signaling has been documented in murine hair follicles and during *Xenopus* gastrulation^59,60^, presenting an additional mechanism of WNT^on^-distal nephron specification in FGF ligand-specific segments. In the proximal tubule, pSMAD1/5 is strongly detected in SSB precursors and Notch activity is required for HNF1B^high^ activation that gives rise to HNF4A^+^ proximal tubule cells^61,62^. Understanding how these different pathways regulate nephron segmentation and how they integrate to form boundaries for proper cell type specification is integral to our goal of robustly constructing an artificial kidney with physiologically functional nephron cell types.

## DISCUSSION

Generating functional iPSC-derived nephrons requires optimized self-organization and identifying external cues that control formation of physiology-imparting cell fates. Here we investigate the nephron axial polarity and signals that tune its development and demonstrate its considerable plasticity. A common approach to generate symmetry breaks along an axis is by modifying developmental pathways (WNT, FGF, BMP, Notch). For instance, WNT and FGF agonists tune axial patterning in embryo models^63,64^, while changes to BMP, WNT, and FGF define midbrain-hindbrain boundaries ^65^. In the nephron, WNT agonist and antagonist have been used in *ex vivo* and *in vitro* studies to control nephron patterning^28,66,67^, while small molecule inhibitors added to kidney organoid cultures can expand specific domains^33,66,68,69^. Our recent work demonstrated that providing signal inputs in the form of ligand-secreting synthetic cellular organizers can locally control axial identities and axial morphogenesis^52^. Here we show that the nephron axis is controlled by WNT, FGF, and BMP signaling, that can be tuned to generate cell identities on demand.

### Axial nephron patterning is regulated by integration of pathways

Given its rapid and reproducible patterning, the nephron tubule is ideal for understanding how signal transduction regulates cell type specification in mammalian development. We adopted a kidney organoid model^29^, developed a protocol to direct differentiation towards distal nephron cell fates, and demonstrate that WNT and FGF pathways cooperatively drive distal nephron identities. Our distal-enriched (DE) protocol differentiated distal axial identities in an imposed transient WNT^ON^ and BMP^OFF^ state that autonomously activated FGF8, and by auto- or paracrine FGF signaling drove early distalization, priming thick ascending loop of Henle-like cell fates. In DE organoids where FGF signaling was blocked (DE + FGF^OFF^), the nephroids reverted their distal axial programming and instead formed proximal tubule and podocyte precursors, fates otherwise not found in DE nephroids. These findings show that FGF signaling acts as a dominant pathway in DE conditions and promotes distal identities by outcompeting or actively suppressing proximal axial fates. Taking this one step further, in DE and FGF^OFF^ organoids where BMP/SMAD signaling is also blocked (i.e., DE and sustained FGF^OFF^ + BMP^OFF^), cells revert to distal axial identities, thus highlighting BMP/SMAD driving proximal cell fates. Our findings illustrate a model where nephron patterning is set up by opposing distalizing WNT/FGF and proximalizing BMP signaling, supporting the hypothesis that FGF and BMP compete to drive nephron fates.

It is not known how these signaling pathways interact to define nephron segment boundaries and specify axial nephron identities. Mouse models show that WNT/β-catenin, BMP, FGF and Notch signaling are all required for nephron patterning^19–22^. We do know that nephrons develop their distal-to-proximal identity against the collecting duct^2,70^ and recent human spatial transcriptomic data shows that WNT/β-catenin target genes are activated in the earliest stages of nephrons in those nephron cells closest to the *WNT9B* expressing collecting duct, and iPSC-derived nephroids respond to canonical WNTs by driving distal axial cell fates^52^. The initial PD symmetry break is therefore likely driven by canonical WNT ligands. At the other end of the early nephron, we here show that cells display active BMP/SMAD signaling (**Figure S3c**), which is absent in distal cells. These two poles set up an early axial asymmetry with distal β-catenin and proximal BMP/SMAD signaling. Consistent with our data dividing the axis into WNT^ON^/BMP^OFF^ and WNT^OFF^/BMP^ON^ poles, are mouse experiments demonstrating that activation of WNT/β-catenin and simultaneous inhibition of BMP/SMAD suppresses podocyte cell fates while driving a Cdh1^+^ distal fate^28^.

A WNT-BMP axial division is reflected in control organoids, which display high pSMAD throughout nephroids and infrequently form distal domains. Shifting these cells to a WNT^ON^/ BMP^OFF^ state between days 11-12, is sufficient to enforce a distal-dominant program leading to cells adopting a distal axial identity. What remains unclear is why mouse nephron cells in WNT^ON^/ BMP^OFF^ states upregulate Tcf/Lef reporters to supraphysiological levels^28^, which would indicate that BMP not only specifies proximal states but also more broadly controls WNT/β-catenin signaling levels.

### FGF signaling as an effector of *β*-catenin-mediated nephron distalization

*FGF8/Fgf8* and *FGFR1*/*Fgfr1* expression patterns appear conserved in developing mouse and human nephrons, suggesting that FGF8 binds to and signals through FGFR1 in the developing mammalian nephron^51^ and in this study. In mouse perturbation experiments, Fgf8 is necessary for the survival of tubular nephron cells and is required for loop of Henle cells to form^51^. This function is underscored by localization of human FGF8 to the medial domain of the late SSB, which may account for the bias of our DE organoids towards loop of Henle/macula densa precursors, and reduction of SLC12A1^+^ tubules upon suppression of FGF signaling (**Figure S7h**). In mice, Fgf8 is a direct target of β-catenin^71^, and it is one of the first genes upregulated in the human PTA, adjacent to the *WNT9B* expressing collecting duct^52^. While a direct WNT-to-FGF connection is likely, it is unclear how FGF8/Fgf8 expression propagates along the nephron axis. However, this proximal moving wave is reminiscent of the proximal wave of Notch ligand, JAG1^3,72^. While Notch signaling is not the focus of our study, Notch is required for the development of nephron cell types^61,73^ and is needed for the Hnf1b^high^ state observed in Jag1^high^ proximal tubule precursors, preceding expression of *HNF4A/Hnf4a* in mouse and human SSBs^33,61^. Further analyses demonstrating the mechanisms by which both FGF and NOTCH signaling shifts proximally, and their interactions with WNT and BMP pathways are therefore required.

## Limitations of Study

The kidney organoid models presented here show an effective and reproducible approach to pathway modulations and axial tuning using small molecule inhibitors. A challenging aspect of this study is characterizing the molecular and cellular targets affected by our pathway changes, which are compounded by the interplay between different pathways. Our model consists of primarily nephron and interstitial cells and the inhibitors can therefore target both populations simultaneously. Developing new approaches such as those described by Huang et al., 2024^74^, where iPSC-derived nephron progenitors are propagated in isolation and then induced to form nephrons, could be a powerful future approach to directly target only nephron cells. However, in *in vivo* environments, the constant crosstalk between the developing nephron and surrounding non-nephron cells is likely critical for normal patterning. While our models provide mechanisms for axial nephron patterning and targeted precursor enrichment, more work is required to delineate the maturation cues necessary for functional nephrons to form *in vitro*.

## Acknowledgements

We thank all past and present members of the Lindström Laboratory and express our gratitude to Dr. Andrew McMahon and Dr. Zhongwei Li for their input. We further thank Dr. Seth Ruffins and the USC Stem Cell Optical Imaging Facility for assistance with microscopy and Dr. Bernadette Masinsin and the Flow Cytometry Facility for help with cell isolation. We thank Kirsten Frieda, Linus Eng, Jay Mehta, and Elizabeth Collins for assistance with spatial transcriptomic work. This work was supported as follows: This work was funded by Lindström lab funding from USC Department of Stem Cell Biology and Regenerative Medicine Startup Fund (NOL), National Institutes of Health under award number NIH/NIDDK R01DK136802 (NOL), American Society of Nephrology/United States Department of Health and Human Services KidneyX (NOL), and T32 Training Grant in Development, Stem Cells, and Regenerative Medicine from NICHD (T32HD060549) (JS).

## Author Contributions

MAA and NOL conceived and executed the project. NOL directed the research and acquired funding. MAA performed experiments. MAA generated organoids, captured, and analyzed data. CCF and RC prepared samples for spatial transcriptomics, JS performed organoid optimization for HNF4A-YFP iPSCs. MAA and NOL prepared and assembled figures. All authors read and provided feedback on the manuscript.

## Declaration of Interests

The authors have filed for intellectual property rights relating to this work.

## SUPPLEMENTAL TABLE LEGENDS

**Table S1. DESeq analyses of total mRNA and single cell sequencing data from DD14 and DD28 control and DE organoids.** This table contains sheets of DESeq-derived differentially expressed genes of control vs DE organoids using data from either total mRNA sequencing or single cell RNA sequencing. The “Readme” sheet contains detailed descriptions of each tab. Related to Figure 1, 2 and S6.

**Table S2. Seurat-based differential gene expression analyses of annotated clusters of DD14 organoid and week 14, 17 human kidney single cell RNA sequencing data.** This table contains sheets of differentially expressed genes of all Seurat objects presented in this study. The “Readme” sheet contains detailed descriptions of each tab. Related to Figure 2, 3, S2 and S5.

## EXPERIMENTAL MODEL AND SUBJECT DETAILS

### Human Kidney Studies

Kidney samples were collected under Institutional Review Board approved protocols (USC-HS-13-0399 and CHLA-14-2211) for previous studies^15,75,76^. Following patient decision for termination, informed consent for donation of the products of conception for research purposes was obtained and samples collected without patient identifiers. The person obtaining informed consent was different than the physician performing the termination procedure, and the decision to donate tissue did not impact the method of termination. Developmental age was determined according to the American College of Obstetrics and Gynecology guidelines^77–79^. Whole intact kidneys were delivered for analysis on ice at 4LC in high glucose DMEM (Gibco) supplemented with 10% fetal bovine serum (GenClone) and 25mM HEPES (Gibco).

### Human Induced Pluripotent Stem Cells (HiPSCs)

iPSCs derived from human foreskin fibroblasts were generously gifted by Dr Amy L. Ryan (University of Iowa). HNF4A-YFP iPSCs (male) were obtained from the ReBuilding a Kidney (RBK) Consortium.

## METHOD DETAILS

### HiPSC-Derived Kidney Organoid Differentiation

iPSCs are cultured in Matrigel (Corning) -coated 12-well plates in Essential 8 medium (Gibco) supplemented with Y-27632 (Tocris) for 24 hours, after which medium is replaced with Essential 8 medium. When the cells are about 60% confluent (∼ 48 hours), iPSCs are passaged and seeded for differentiation into kidney organoids using a modified Takasato protocol^80^. Briefly, on DD0, iPSCs are seeded at a density of 17.5k cells/well in a Matrigel-coated 12-well plate in Essential 8 medium supplemented with 5μM Y-27632. After 6 hours, 9 μM CHIR99021 (Tocris) in TeSR-E6 (STEMCELL Technologies) is added and replenished every other day for 5 days, followed by TeSR-E6 containing 200 ng/ml FGF9 (Bio-techne) and 100 ng/ml Heparin (Sigma-Aldrich) between DD5 and DD7. On DD7, cell aggregation is achieved by adding 30 μl of 100,000 cells in TeSR-E6 containing 10 μM Y-27632 into a round-bottom low-attachment 96-well plate. The plate is spun at 100 ×g for 2 minutes in a swing bucket centrifuge and incubated at 37LC for 2 hours. The aggregates are then transferred into 6-well Transwell plates (STEMCELL Technologies), following the Takasato protocol. For our distal-enrichment protocol, 10 μM LY294002 (Tocris) is added to the TeSR-E6 medium supplemented with FGF9 and Heparin from DD10-11. On DD11, TeSR-E6 medium containing FGF9 and Heparin is supplemented with 10 μM LY294002, 1.5 μM CHIR99021 and 1μM LDN 193189 dihydrochloride (Tocris) and added to these organoids. On DD12, medium is changed to TeSR-E6 with no growth factors for both control and treated groups. Organoids are cultured until DD28.

For validation experiments of PI3K inhibition, 10 μM LY29 is replaced with 0.75 μM GDC-0941 (Selleck) from DD10-DD12, with 1.5 μM CHIR and 1 μM LDN added to this media from DD11-DD12. For our BMP inhibition validation in our DE protocol, organoids were treated with 10 μM LY29 (DD10-DD12), to which 1.5 μM CHIR and 2.5 μM Dorsomorphin (Sigma-Aldrich) were added from DD11-DD12. For our FGFR1 inhibition experiments, 1 µM SU-5402 (Sigma-Aldrich) is added from DD11-DD12 to TeSR-E6 supplemented with FGF9, Heparin, LY29, CHIR and LDN in the amounts specified above, then TeSR-E6 with no growth factors from DD12-DD14. HNF4A-YFP iPSCs^48^ were differentiated with the same differentiation protocol described above, with the exception of DD0-DD5, where they were treated with 7 μM CHIR99021, based on the differentiation protocol outlined in Schnell et al., 2024^33^.

### Immunofluorescence Analysis

#### Immunostaining of Human Kidney Cryosections

Frozen human week 18.6 kidney cryosections were thawed at room temperature (RT) for approximately 5 minutes. Antigen retrieval was performed using 1X Citrate buffer (Sigma-Aldrich) on high pressure for 1 minute in a pressure cooker. The slides were rinsed with deionized water and incubated for 1 hour in blocking buffer solution (1.5% SEA BLOCK blocking buffer (Thermo Scientific) + 0.1% Triton X-100 (MD Millipore) in 1X PBS (Thermo Scientific)). The slides were incubated overnight at 4LC in primary antibodies diluted in blocking buffer solution (see “*Immunostaining of Whole Kidney Organoids*” section below for dilutions used), rinsed in PBS-T (0.1% Triton X-100 in 1X PBS) and incubated in secondary antibodies diluted in blocking buffer solution for 1 hour at room temperature; secondary antibodies conjugated with Alexa Fluor 488, 555, 594, and 647 were all diluted to 1:500. Once rinsed, the slides were incubated in 1 μg/ml Hoechst 33342 (Thermo Scientific) in PBS-T for 5 minutes at RT for nuclei staining, followed by a final rinse in 1X PBS. Sections were then mounted with Immu Mount (Thermo Scientific).

#### Immunostaining of Whole Kidney Organoids

Whole organoids were fixed with 4% paraformaldehyde (Electron Microscopy Sciences) in 1X PBS – 1.2 ml of 4% paraformaldehyde was added under the transwell and incubated for 10 minutes on ice, after which an additional 1 ml of 4% paraformaldehyde was added to the top of the organoids and incubated for 10 minutes on ice. After incubation, the 4% paraformaldehyde was replaced with cold 1X PBS. Fixed organoids were cut out of the transwells and transferred into blocking buffer solution. The organoids were incubated on a nutator at 4LC for 1 hour, after which they were incubated overnight on a nutator at 4LC in primary antibodies diluted in blocking buffer solution. The primary antibodies used were as follows: Laminin β1 (Santa Cruz Biotechnology, sc-33709, 1:250), HNF4A (R&D Systems, MAB4605, 1:200), POU3F3 (Thermo Scientific, PA564311, 1:100), Mafb (Novus (R&D), MAB3810, 1:500), GATA3 (Novus (R&D), AF2605, 1:200), Jag1 (Novus (R&D), AF599, 1:100), SLC12A1 (Sigma-Aldrich, HPA018107, 1:200), PAX8 (abcam, ab189249, 1:100), WT1 (abcam, ab89901, 1:500), LEF1 (Cell signaling, 2286S, 1:200), CDH1 (BD Laboratories, 610181, 1:200), PAPPA2 (R&D Systems, AF1668, 1:100), HNF1B (Thermo Scientific, MA5-24605, 1:500), Pax2 (Thermo Scientific, 716000, 1:100), TFAP2A (Santa Cruz Biotechnology, sc-12726, 1:200), IRX1 (Sigma-Aldrich, HPA043160, 1:200), SIX1 (Cell signaling, 12891S, 1:300), Pax2 (R&D Systems, AF3364, 1:50), TMEM52B (Novus (R&D), NBP2-49272, 1:100), MEIS1/2/3 (Active Motif, 39096, 1:1000), ZO-1 (Thermo Scientific, 33-9100, 1:200), PODXL (R&D Systems, AF1658, 1:300), Renin (R&D Systems, AF4090, 1:100), MECOM (R&D Systems, MAB75061, 1:100), hErbB4 (R&D Systems, MAB1131, 1:100), LEF1 (Santa Cruz Biotechnology, sc-374412, 1:200), CD31 (abcam, ab9498, 1:250), ESRRγ (Sigma-Aldrich, HPA044678, 1:100), GATA3 (Cell Signaling Technology, 5852T, 1:100), and acetyl-alpha tubulin (Sigma-Aldrich, MABT868, 1:500). After a 3x wash in PBS-T (1 hr each), organoids were incubated overnight on a nutator at 4LC in secondary antibodies diluted in blocking buffer solution. Secondary antibodies conjugated with Alexa Fluor 488, 555, 594, and 647 were all diluted to 1:500. Following incubation, the organoids were washed 3x in PBS-T (1 hr each), followed by a 35-minute incubation on a nutator at 4LC in 1 μg/ml Hoechst 33342 in PBS-T. After a final rinse in 1X PBS, organoids were serially dehydrated in 50%, 75% and 100% methanol in 1X PBS for 30 minutes each, then incubated at RT in 50% methanol in a BABB solution (benzyl alcohol (Sigma-Aldrich) and benzyl benzoate (Sigma-Aldrich) in a 1:2 ratio). The organoids were then fully cleared in 100% BABB and stored at 4LC.

### Immunohistochemistry Analysis

#### Paraffin Embedding and Sectioning

Week 16 human kidneys were fixed in 4% paraformaldehyde in 1X PBS for 1 hour at 4LC and washed 2x in 1X PBS for 5 minutes each. They were transferred to cassettes and stored in 70% ethanol in 4LC and transferred to the USC Translational Research lab histology core for paraffin processing and sectioning in 5-μm sections. DD14 kidney organoids were fixed in 4% paraformaldehyde following the same protocol described in the “*Immunofluorescence Analysis*” section. The paraffin embedding protocol for organoids was adapted from Wörsdörfer et al., 2020 ^81^. Briefly, fixed organoids were detached from Transwell membranes and embedded in 300 ul of 1% agarose (Avantor) in 1X PBS at 60LC in 10 mm x 10 mm x 5 mm cryomolds (Tissue Tek, Cat# 4565) and allowed to set. The agarose blocks were transferred to tissue cassettes (Simport, Cat# M511-10) and stored in 70% ethanol in 4LC overnight. Paraffin processing was done using the Tissue Tek VIP tissue processor (Sakura) and the agarose blocks were embedded into a paraffin block. The organoid blocks were sectioned into 5 μm-thick sections using a sliding microtome and dried overnight at 60LC in a lab oven.

#### Antibody Staining and Hematoxylin Counterstaining

Paraffin sections were dewaxed with the following solution series: 100% Xylene (Sigma-Aldrich) for 10 minutes (2x), 100% ethanol for 1 minutes, 95% ethanol for 1 minute, 80% ethanol for 1 minute, 60% ethanol for 1 minute, 30% ethanol for 1 minute and 2x rinse in deionized water. Antigen retrieval was performed as described in the “*Immunofluorescence Analysis*” section, and the sections were incubated in 0.1% TBS-T (0.1% Tween-20 in 1X TBS (Cell Signaling Technology)) + 1.5% SEA BLOCK blocking buffer for 1 hour at RT. Sections were incubated in pSMAD1/5 (Cell Signaling, 9516T, 1:100) diluted in 0.1% TBS-T + 1.5% SEA BLOCK blocking solution overnight at 4LC. The slides were washed 2x in 0.1% TBS-T for 5 minutes each. Endogenous peroxidase activity was blocked with 3% hydrogen peroxide (Sigma-Aldrich, Cat# 216763) for 10 minutes at RT and rinsed 2x with deionized water. The slides were incubated in biotinylated secondary antibody in 0.1% TBS-T + 1.5% SEA BLOCK blocking solution for 30 minutes, then washed 3x with 0.1% TBS-T for 5 minutes each. An avidin-biotin solution was prepared from the VECTASTAIN ABC kits (Vector laboratories) using manufacturer instructions to visualize biotinylated secondary antibodies and slides were incubated in ABC solution for 30 minutes at RT. Slides were washed again 3x with 0.1% TBS-T for 5 minutes each. The slides were developed using the ImmPACT DAB substrate kit (Vector laboratories) for 30 minutes and development was stopped by plunging the slide into deionized water. The slides were counterstained with hematoxylin (Sigma-Aldrich) for 10 seconds, rinsed with deionized water and mounted using Immu Mount.

#### Image Acquisition and Analysis

Fluorescent immunostains were imaged on the Leica SP8 with DLSM (Leica) and Stellaris 8 (Leica) confocal microscopes. Z-stack images were acquired at 3μm intervals. Image processing for images taken on Leica microscopes was done on LAS X image analysis software (Leica). The immunohistochemistry images were taken on the ZEISS Axioscan 7 (ZEISS) and processed on ZEISS ZEN lite image analysis software (ZEISS).

#### Spatial Genomics Analysis on Human Kidney Samples

Week 16 human kidneys embedded in OCT (Electron Microscopy Sciences) were cryosectioned at 10μm thickness and placed on coverslips provided by Spatial Genomics. They were fixed in 4% paraformaldehyde in PBS for 15 minutes at RT. Sections were washed 4 times in 1x PBS for 5 minutes at RT and dehydrated in 100% isopropanol for 30 seconds at RT. The sections were air dried, stored at −80LC and shipped to Spatial Genomics, Inc in dry ice for sequencing. Select probes consisted of 250 cell-state and/or lineage-defining genes that represented all 5 nephron cell types. These genes were derived from differential gene expression analysis of human single-nuclei RNA sequencing data. Data collection and imaging were performed by Spatial Genomics and visualization was done using the Spatial Genomics Viewer software, with screengrabs obtained at 0.32x zoom.

### Bulk mRNA Sequencing and Analyses of Human Kidney Organoids

#### Sample Collection and Preparation

All bulk RNA samples were collected in duplicate (2 biological replicates). Kidney organoids were collected at select timepoints (2-3 organoids pooled per condition) into a 1:100 solution of lysis buffer RLT (Qiagen) and β-Mercaptoethanol (Sigma-Aldrich) and homogenized by pipetting for 30 seconds. RNA was isolated from samples using the Qiagen RNeasy kit for RNA purification (Qiagen) and total RNA was quantified on a NanoDrop spectrophotomer (Thermo Scientific). mRNA from the samples was sequenced by Novogene using the Illumina NovoSeq X Plus PE153 platform (Illumina).

#### Data Processing

Raw FASTQ files were aligned to the human genome using STAR alignment^82^ and feature counts were generated from BAM files using the *featureCounts* function of the Rsubread package^83^. TPM normalization was performed, and genes were annotated on Partek Flow, where .txt files were generated with TPM values for each gene in each sample. For generating heatmaps of bulk mRNA sequencing data, a matrix of TPM values was loaded on R, and mitochondrial and ribosomal genes and gene rows with TPM values of <10 were excluded. The *pheatmap* function of the pheatmap package was used to generate heatmaps that used hierarchical clustering. The *DESeqDataSetFromMatrix* function from the DESeq2 package^84^ was used to perform DESeq analysis on paired samples and *EnhancedVolcano* function generated volcano plots, where a p-value cutoff of 0.05 and log_2_ fold change cutoff of 1.5 was set.

### Single-cell RNA Sequencing and Analyses of Human Kidney Organoids

#### Sample Collection and Preparation

Kidney organoids were collected at select time-points (9 organoids each for DD14 control and DE organoids) and dissociated into single cells using Accumax (Stemcell) for 45 minutes at 37 LC, with the organoids pipetted at 5-minute intervals with a P1000 tip 10 times. Once the cells were fully dissociated, the Accumax enzyme was neutralized using twice the volume of cold AutoMacs buffer (Miltenyi). The tubes were centrifuged at 300 ×g for 3 minutes at 4LC, the supernatant was aspirated, and the dissociated cells were resuspended in 2 ml of cold AutoMacs buffer. The cells were strained using a 40μm cell strainer and chased with 1 ml of cold AutoMacs buffer, then centrifuged again at 300 ×g for 3 minutes at 4LC. The supernatant was aspirated, and the cells were resuspended with 500 μl of cold AutoMacs buffer containing nuclear dyes DRAQ5 (1:1000) and DAPI (1:1000) and FAC-sorted for live cells on the ARIA II FACS (BD Biosciences). The resuspended cells were transferred to round-bottom 15-ml tubes. Using dissociated cells resuspended in AutoMacs buffer as a negative control for gating, the dissociated cells resuspended in the nuclear dyes were sorted for single-live cells (DRAQ5^+^, DAPI^-^) for collection. These cells were then fixed using the Parse Biosciences WT v2 kit and protocol and for each datapoint, 7 sublibraries were prepared for SPLiT-seq and sequenced separately by Novogene using the Illumina NovaSeq X Plus 25B PE150 platform (Illumina) with a read depth of ∼63k reads/cell. Raw FASTQ files of each sublibrary were demultiplexed and aligned to the human genome using split-pipe from Parse Biosciences, where corresponding conditions were assigned to each sublibrary.

#### Seurat Object Processing

Seurat objects were made for each sublibrary using the Seurat package (V4) in R^85^ and initial quality control metrics were run to cutoff low quality cells (1000-8000 genes per cell, percent mitochondria < 20). For each condition, the 7 Seurat objects (1 object per sublibrary) were merged using *merge* function and no batch effects were observed. However, cycling genes were significantly contributing to cell heterogeneity as cells separated by cell cycling phases, thus we log-normalized the merged objects and regressed out cell cycle scores^86^ while scaling features in our datasets using the *ScaleData* function. Non-linear dimensionality reduction was performed using *RunUMAP* and *FindNeighbors* using 30 principal components. The objects were clustered at a resolution of 1.5. For objects from each condition, differential gene expression analysis was performed on log-normalized transcripts using *FindAllMarkers* function, where only genes detected in a minimum of 25% of cells in either population were tested and the cutoff for log fold change was 0.25. From the control organoid Seurat object, cells from clusters 21, 25, 15, 17, 6, 24, and 12 were identified as interstitial and subset out to retain nephron-only clusters. Similarly, from the DE organoid Seurat object, clusters 24, 4, 19, 18, 12, and 20 were subset out. In each instance, Nephron-only subset objects were reclustered at a resolution of 1.5.

Nephron-only objects from control and DE organoids were merged using the *merge* function, where cell cycle regression was performed again during scaling. Differential gene expression analysis was performed on log-normalized transcripts using the same criteria described above. DESeq-derived differential gene expression between control and DE nephron cells was determined using the *FindMarkers* function and the results were plotted on a volcano plot using the *EnhancedVolcano* function, where a p-value cutoff of 0.5 and log_2_ fold change cutoff of 1.5 was set.

#### Reanalysis of Published Single-cell RNA Sequencing Data

Single cell RNA sequencing data of week 14^49^ and week 17^1^ human kidney were downloaded from the Gene Expression Omnibus (accession numbers: GSE139280, GSE127344). The Seurat package (v3) was used for quality control analyses, SCTransform data integration^87^ of the 2 datasets. Based on differential gene expression of known kidney cell type markers, clusters were annotated with corresponding cell types and differential gene expression analysis was performed using *FindAllMarkers*.

#### Transcription Factor Analysis

For analysis of distal nephron-enriched transcription regulators, we examined gene expression levels of transcription factors from the human^88^ and mouse^89^ transcription factor atlases – the mouse atlas was converted into its human ortholog using Ensembl BioMart ^90^ as described in Kim, S. et al., 2024^6^. The enrichment of transcription factor detection in TFAP2A^+^ clusters was shortlisted by feature plot visualization and represented in **Figure S5**.

##### Multimodal Reference Mapping of Human and Organoid Single-cell RNA Sequencing Data

The annotated clusters of the single cell human kidney Seurat object was used to predict cell types in the DD14 control and DE nephroid datasets following the Seurat v4 reference mapping vignette developed by the Satija Lab^91^. Briefly, the merged organoid dataset was split by condition and treated as separate query samples for finding anchors to map them to the human kidney reference. The *ggplot* package was used for visualization of the number of cells in each condition that were represented in the predicted cell types.

##### QUANTIFICATION AND STATISTICAL ANALYSIS

Quantification of symmetric and asymmetric nephroids was performed on ImageJ (Fiji) using the Cell Counter plugin. Statistical analysis shown on all bar plots was performed using unpaired parametric t-test (GraphPad Prism) for biological replicates (n = 2), where the p-value cut-off is 0.05.

### RESOURCE AVAILABILITY

#### Lead contact

Further information and requests for resources and reagents should be directed to and will be fulfilled by the lead contact, Nils O. LindstrLm.

#### Materials availability

This study did not generate new unique reagents.

---

**Supplemental Figure 1.**
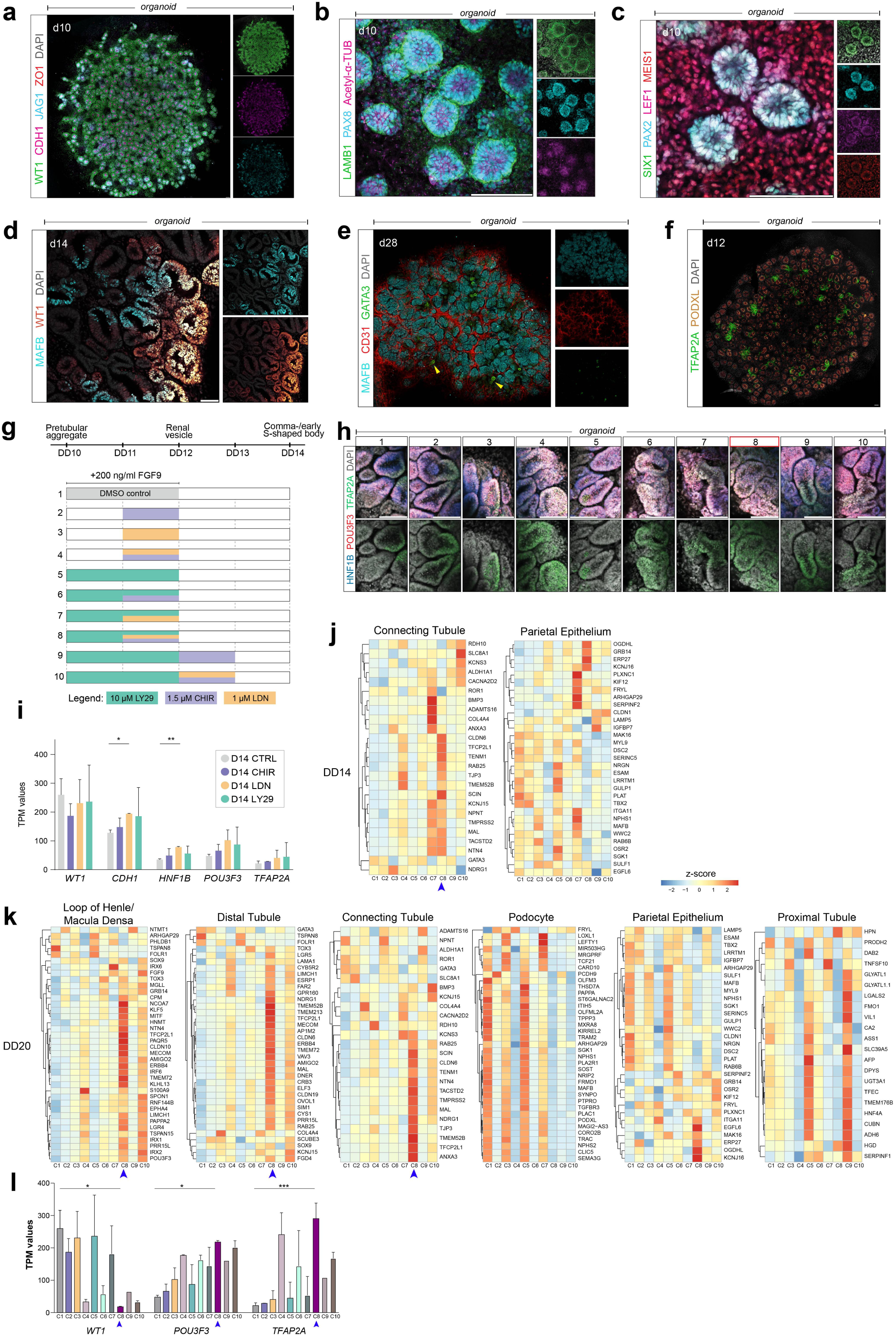
Characterization of developmental stages of kidney organoid differentiation. Related to Figure 1. (a-c) Immunofluorescence stains of DD10 organoids illustrating expression of select induced nephron progenitor markers. Scale bar: 50 microns. (d) Immunofluorescence stain of DD14 control organoids illustrating emergence of MAFB^+^ podocyte precursors. (e) Immunofluorescence stain of DD28 control organoids showing MAFB^+^ glomerular-like structures and surrounding vasculature-like structures and sporadic GATA3^+^ tubules (yellow arrows). Scale bar: 50 microns. (f) Immunofluorescence stain of DD12 control organoid tile scan showing TFAP2A emergence. Scale bar: 50 microns. (g) Schematic of SMI treatment screening to optimize nephron distalization. Top bar shows timeline for SMI treatment for each condition. (h) Immunofluorescence stains of DD14 organoids generated from the 10 condition-screen showing distal transcription factor. Red box denotes condition 8. Scale bars: 50 microns. (i) Bar graph of TPM values show expression levels of *WT1, CDH1, HNF1B, POU3F3* and *TFAP2A* in DD14 organoids (n = 2) treated with different SMIs. Error bars show standard deviation, and top bars show statistical significance between each SMI treatment against control based on non-parametric t test (* - p<0.05, ** - p<0.01). (j-k) Heatmaps of TPM value z-scores of nephron cell type markers at the SSB stage, expressed across 10 differentiation conditions in (j) DD14 and (k) DD20 organoids (n = 2). (l) Bar graph of TPM values show expression levels of *WT1, POU3F3* and *TFAP2A* in DD14 organoids (n = 2) from all 10 conditions. Data are represented as mean ± SD with statistical significance based on non-parametric t test between condition 8 and DMSO control (* - p<0.05, *** - p<0.001).

**Supplemental Figure 2.**
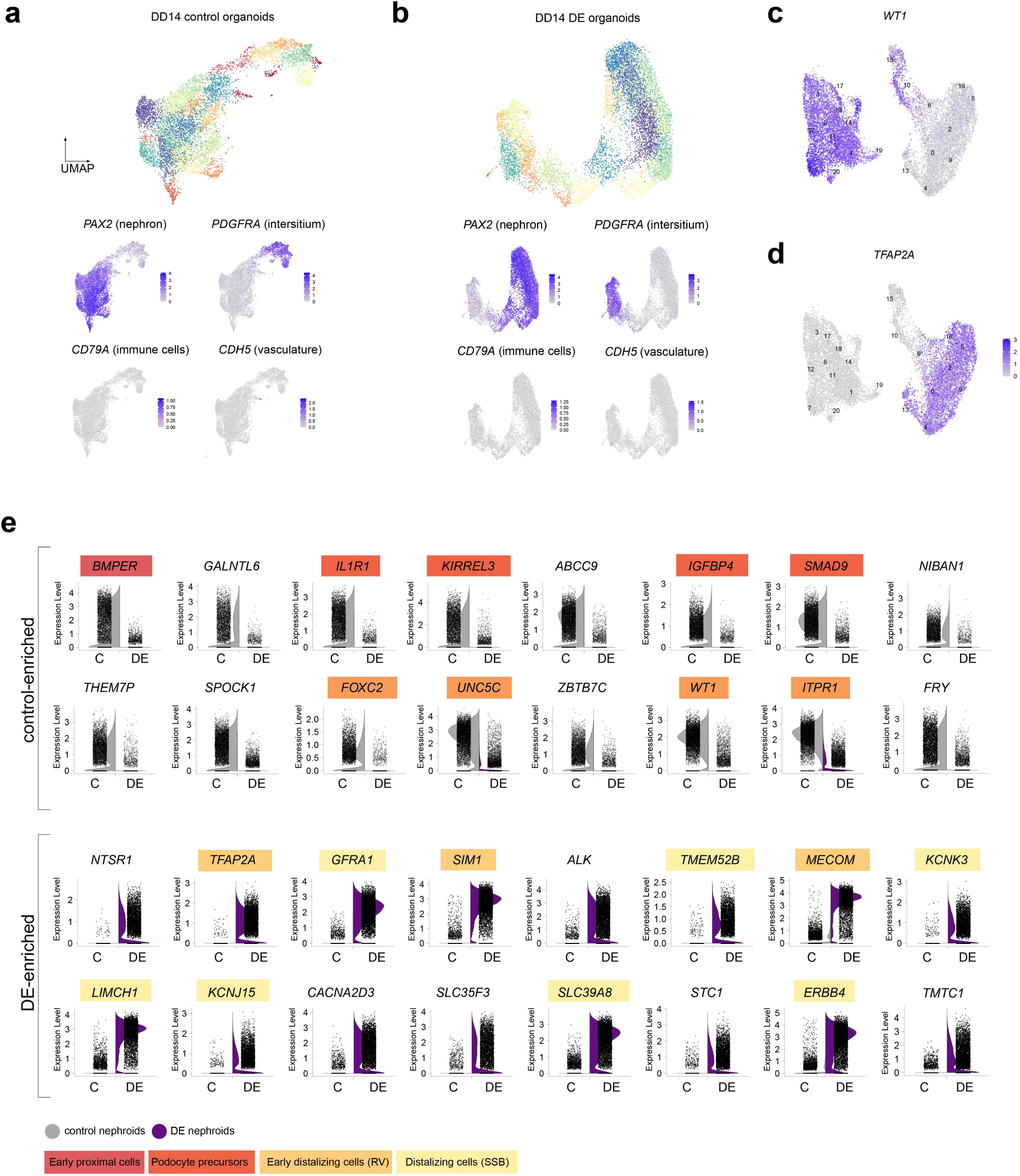
Single cell RNA sequencing of DD14 organoids. Related to Figure 2. (a-b) (Top) UMAPs of single cell sequencing data of DD14 control and DE organoids. (Bottom) Feature plots of select cell type markers in DD14 control and DE organoids. (c-d) Feature plots of DD14 organoid scRNAseq data showing detection of *WT1* and *TFAP2A*. (e) Violin plots split by condition showing total expression of top 16 DEGs in DD14 control (top 2 rows) and DE (bottom 2 rows) organoids from scRNAseq dataset. Colors behind gene names correspond to cell types indicated in Figure 2b.

**Supplemental Figure 3.**
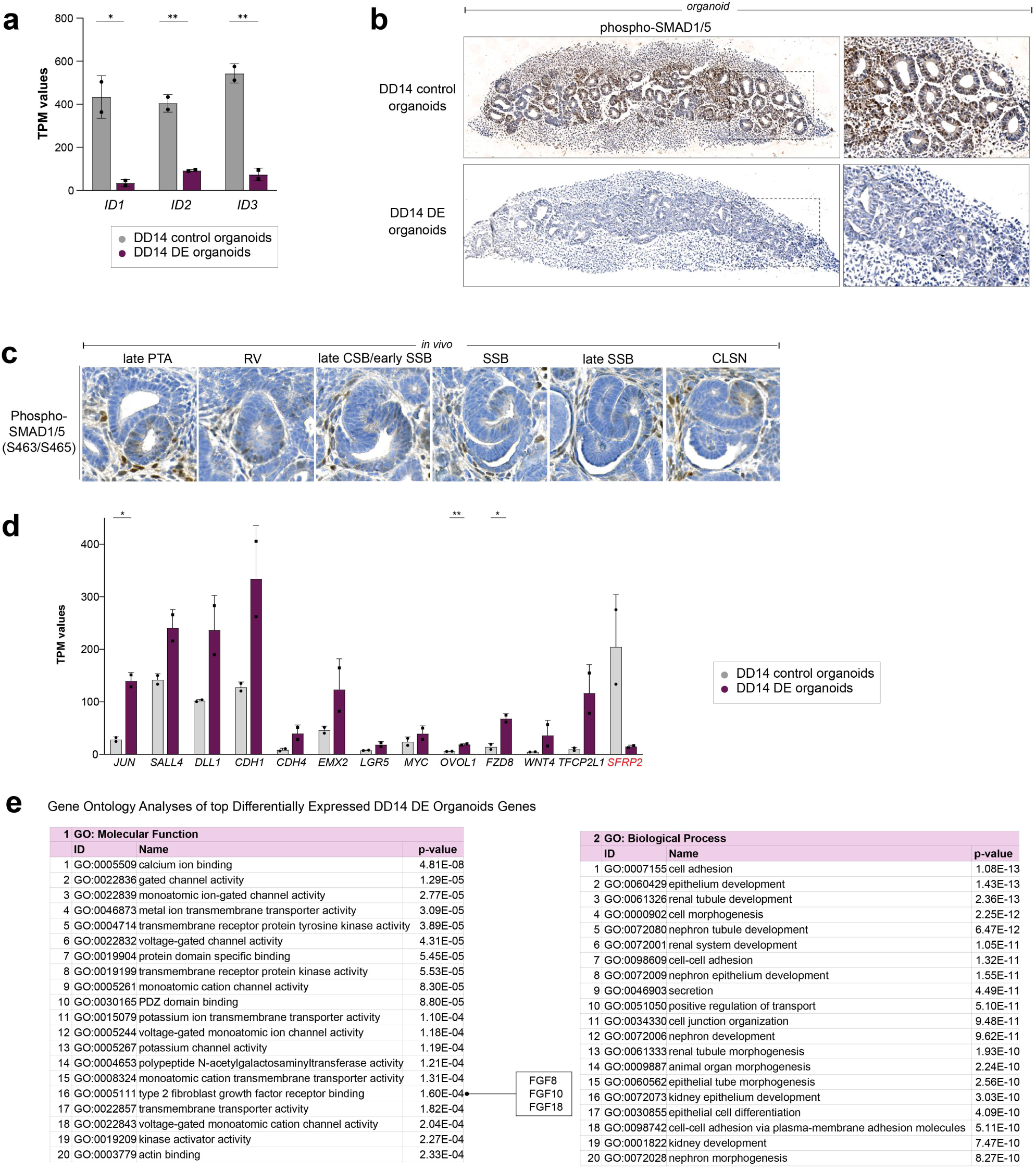
Combined β-catenin activation and BMP inhibition is sufficient to bias organoids towards a distal fate. Related to Figure 2. (a) Bar graph of TPM values of DD14 control and DE organoids of BMP target genes. Error bars show standard deviation, and horizontal bars show statistical significance based on non-parametric t test (* - p < 0 .05, ** - p < 0.01). (b) (top) DD14 control and (bottom) DE organoid cross-sections showing nuclear phospho-SMAD1/5 detection. (c) IHC staining of week 16 human kidneys showing nuclear dual phospho-SMAD1/4 detection. (d) Bar graph of TPM values of DD14 control and DE organoids of canonical WNT target genes. Data are represented as mean ± SD with statistical significance based on non-parametric t test (* - p < 0 .05, ** - p < 0.01). (e) GO term analyses of molecular function and biological process predicted from top DEGs of DD14 DE organoids (log_2_FC > 1.5).

**Supplemental Figure 4.**
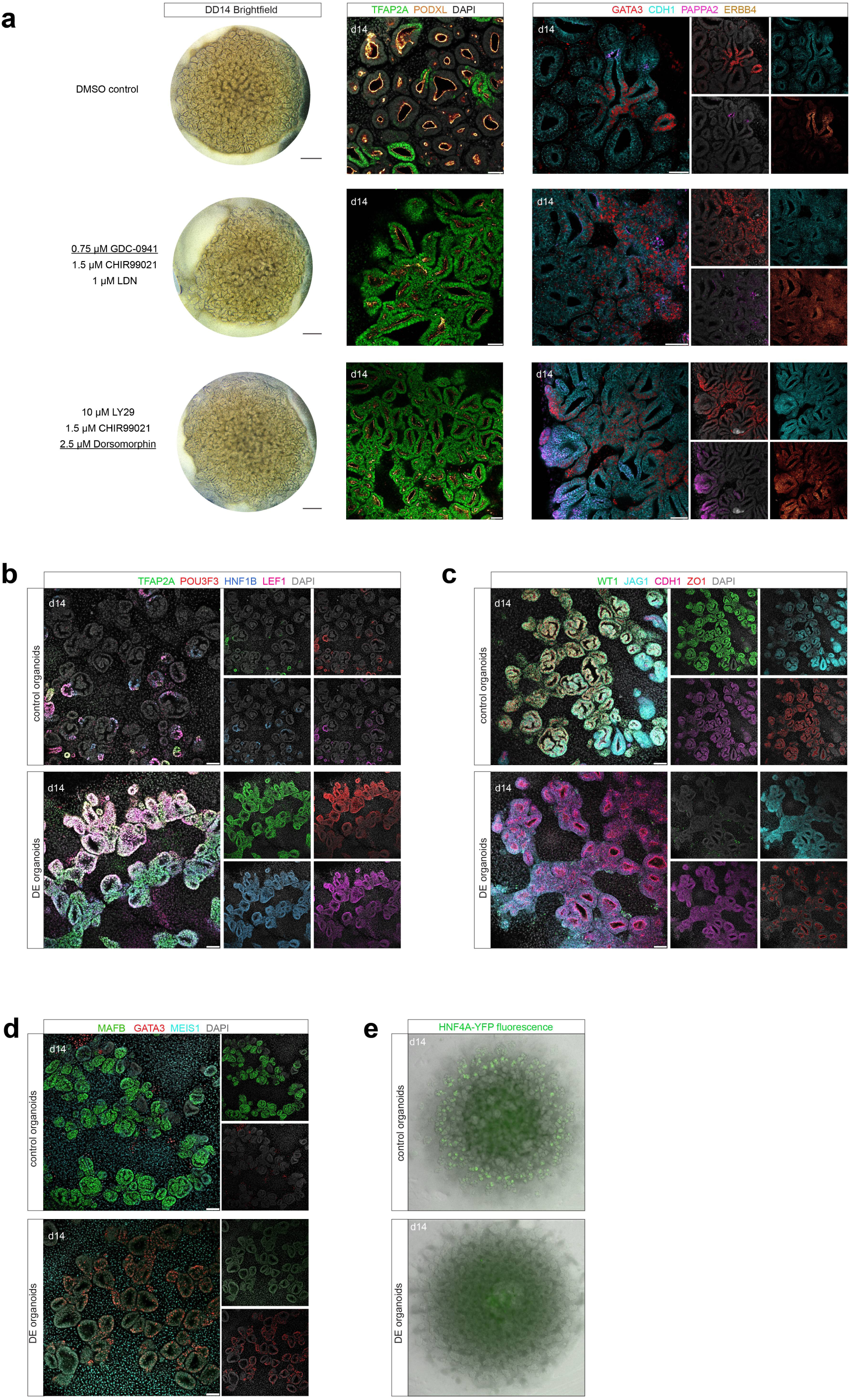
Secondary validation of distal-enrichment protocol. Related to Figure 2. (a) Brightfield images and immunofluorescence stains on DD14 control and organoids treated with dosages indicated (left) showing detection of select distal nephron markers. (b-d) Immunofluorescence stains on DD14 control and DE protocol-treated organoids cultured from HNF4A-YFP iPSCs. Immunostains show detection of select podocyte and distal nephron markers. Scale bar: 50 microns. (e) Live imaging of YFP fluorescence in DD14 control and DE protocol-treated organoids cultured from HNF4A-YFP iPSCs.

**Supplemental Figure 5.**
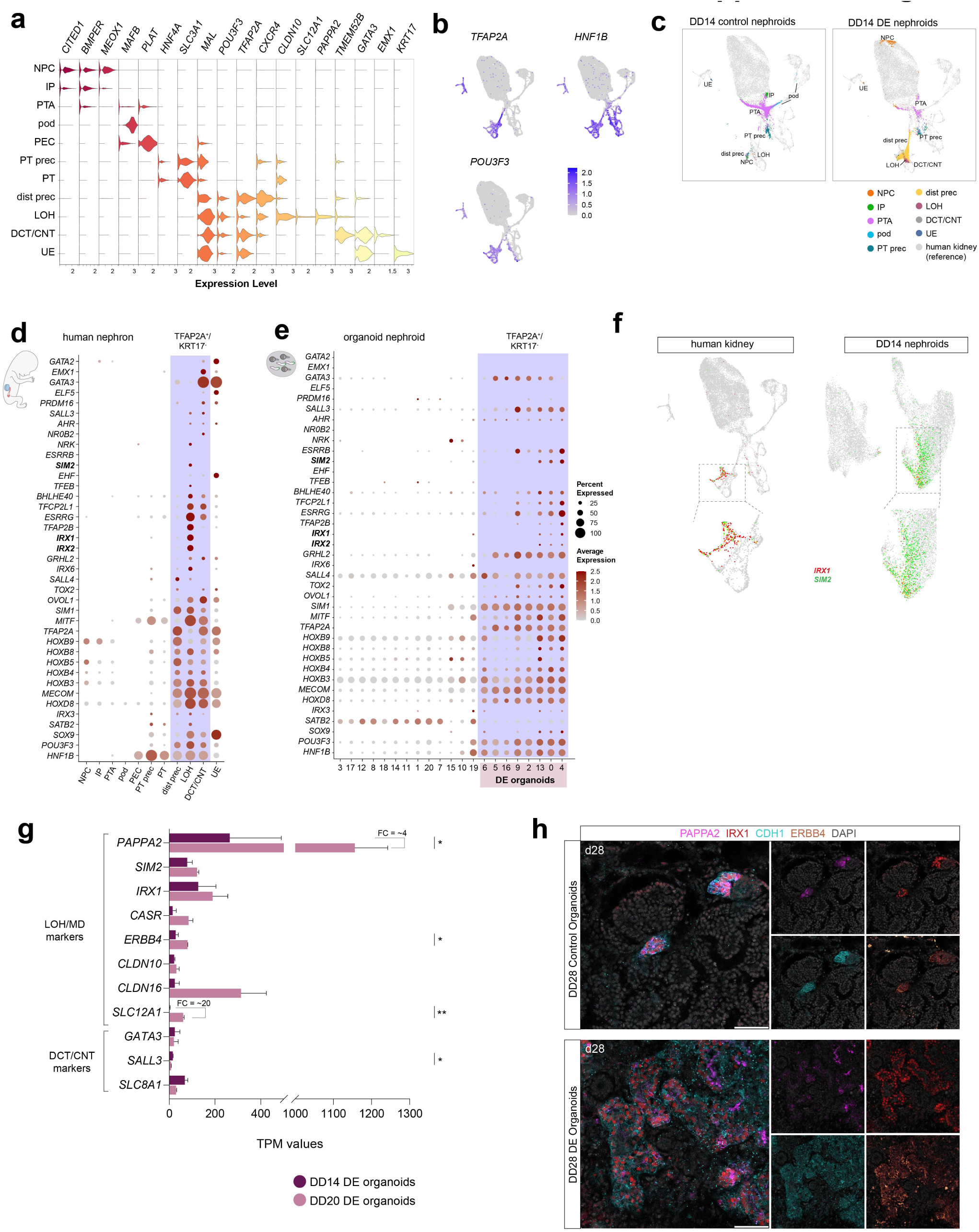
Transcriptional comparison between human and organoid nephron development. Related to Figure 3. (a) Violin plots representing nephron cell type markers used to annotate cell types of single cell RNA sequencing performed on week 14, week 17 human kidney. (b) Feature plots of TFAP2A, HNF1B and POU3F3 detection in human kidney dataset. (c) Annotated mapping of mapped control (left) and DE (right) organoid clusters using the developing human kidney dataset as a reference. (d-e) Dot plot of distal-enriched transcription factor expression in (d) human and (e) organoid scRNA sequencing datasets. Purple box marks nephrogenic *TFAP2A*^+^ clusters. (f) Feature plots showing co-expression of transcription factors *IRX1* (red) and *SIM2* (green) in distalizing cells of human kidney and organoid datasets. (g) Horizontal bar plot showing TPM values of select distal nephron markers in DD14 and DD20 DE organoids. Data are represented as mean ± SD with statistical significance based on non-parametric t test (* - p < 0.05, ** - p < 0.01). FC = fold change. (h) Immunofluorescence stain of DD28 control and DE organoids showing detection of loop of Henle/macula densa precursor markers. Scale bar: 50 microns.

**Supplemental Figure 6.**
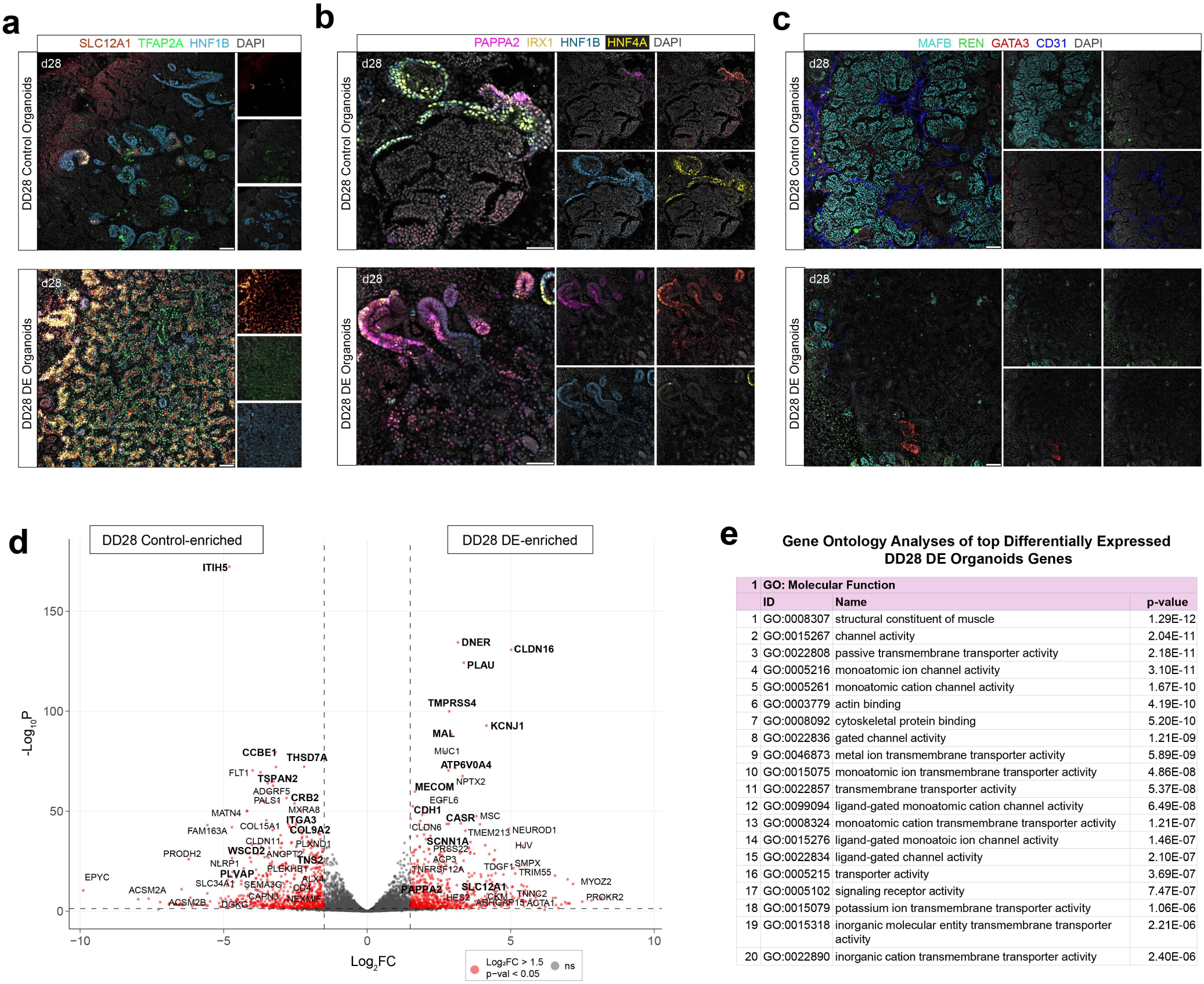
Characterization of DD28 kidney organoids. Related to Figure 3. (a-c) Immunofluorescent stains of DD28 control and DE organoids showing detection of glomerular, proximal tubule, loop of Henle/macula densa and DCT precursor markers. Scale bar: 50 microns. (d) Volcano plot showing DESeq analysis of total mRNA sequencing data of DD28 control versus DE organoids. Dotted lines show 0.05 p-value cutoff (horizontal) and ±1.5-fold change cutoff (vertical). (e) GO term analyses of molecular function and biological process predicted from top DEGs of DD28 DE organoids (log_2_FC > 1.5).

**Supplemental Figure 7.**
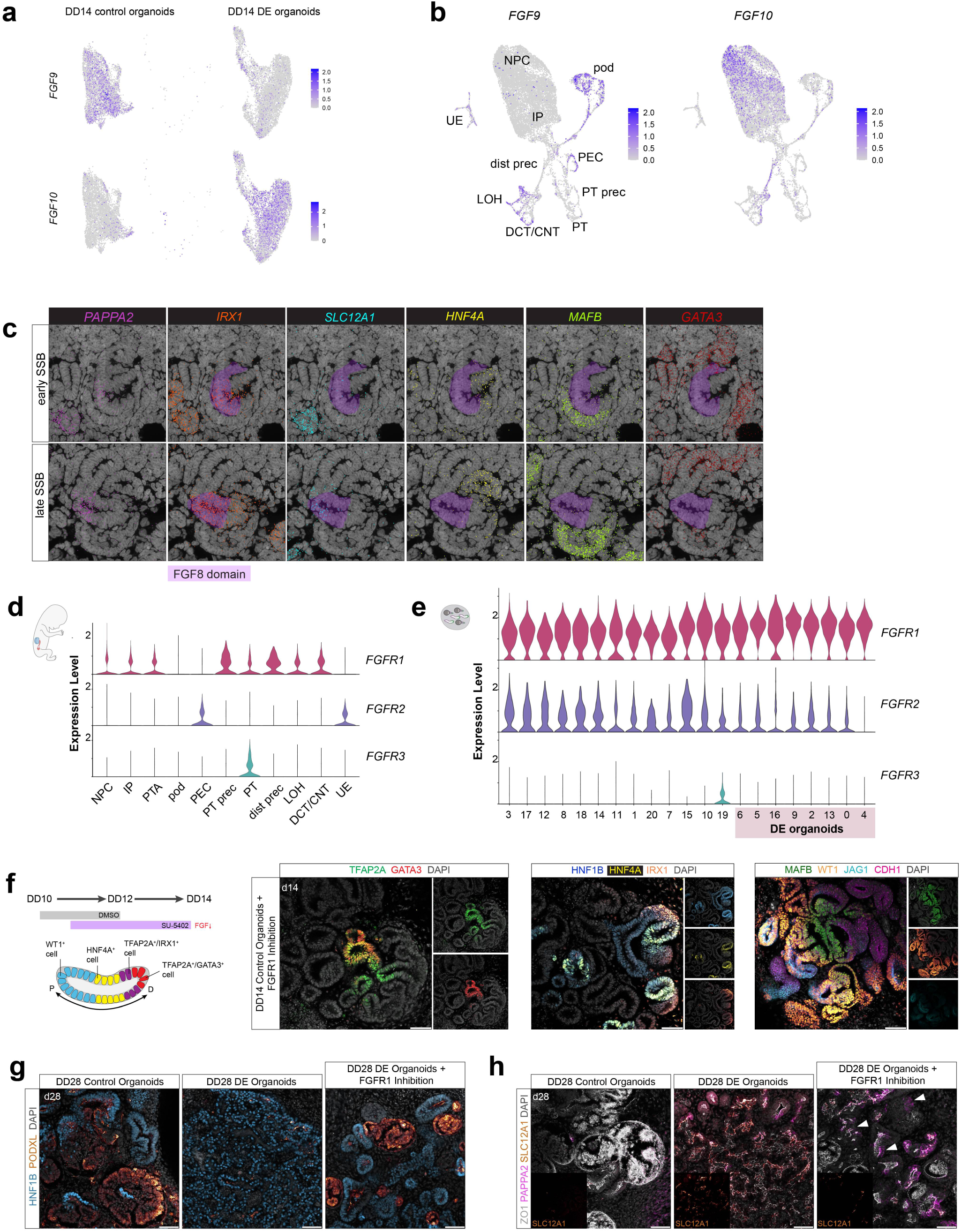
Examining FGF signaling in early nephron patterning. Related to Figure 4. (a) Feature plots of *FGF9* and *FGF10* expression in the DD14 organoid Seurat object, split by condition. (b) Feature plots of *FGF9* and *FGF10* expression in the developing human kidney Seurat object. (c) SeqFISH data of week 16 human kidney showing detection of select markers in early and late SSB. (d-e) Violin plots of FGF receptor expression in each cell type of the (d) human kidney Seurat object and (e) DD14 kidney organoid object. (f) Schematics of SMI treatment and nephron segmentation in DD14 organoids (left) and immunofluorescent stains showing detection of DCT/CNT, proximal tubule, loop of Henle/macula densa and podocyte precursor markers (right). Scale bar: 50 microns. (g-h) Immunofluorescent stains of DD28 control, DE, and DE+FGF-deficient organoids showing (g) podocyte-to-tubule distribution and (h) formation of loop of Henle/macula densa-like precursors.

## Notes

### Competing Interest Statement

The authors (MAA and NOL) have filed for intellectual property rights relating to this work.

## REFERENCES

1. Lindström, N. O. et al. Spatial transcriptional mapping of the human nephrogenic program. Dev. Cell 56, 2381–2398.e6 (2021).

2. Georgas, K. et al. Analysis of early nephron patterning reveals a role for distal RV proliferation in fusion to the ureteric tip via a cap mesenchyme-derived connecting segment. Dev. Biol. 332, 273–286 (2009).

3. Lindström, N. O. et al. Progressive Recruitment of Mesenchymal Progenitors Reveals a Time-Dependent Process of Cell Fate Acquisition in Mouse and Human Nephrogenesis. Dev. Cell 45, 651–660.e4 (2018).

4. Ransick, A. et al. Single-Cell Profiling Reveals Sex, Lineage, and Regional Diversity in the Mouse Kidney. Dev. Cell 51, 399–413.e7 (2019).

5. Lindström, N. et al. Spatial Transcriptional Mapping of the Human Nephrogenic Program. (2020) doi:10.1101/2020.04.27.060749.

6. Kim, S. et al. Comparative single-cell analyses identify shared and divergent features of human and mouse kidney development. Dev. Cell 2023.05.16.540880 (2024) doi:10.1016/j.devcel.2024.07.013.

7. Chen, L., Chou, C.-L. & Knepper, M. A. Targeted Single-Cell RNA-seq Identifies Minority Cell Types of Kidney Distal Nephron. J. Am. Soc. Nephrol. 32, 886–896 (2021).

8. Kobayashi, A. et al. Six2 Defines and Regulates a Multipotent Self-Renewing Nephron Progenitor Population throughout Mammalian Kidney Development. Cell Stem Cell 3, 169–181 (2008).

9. Nishinakamura, R. Human kidney organoids: progress and remaining challenges. Nat. Rev. Nephrol. 15, 613–624 (2019).

10. Wu, H. et al. Comparative Analysis and Refinement of Human PSC-Derived Kidney Organoid Differentiation with Single-Cell Transcriptomics. Cell Stem Cell 23, 869–881.e8 (2018).

11. Tung, C. W., Hsu, Y. C., Shih, Y. H., Chang, P. J. & Lin, C. L. Glomerular mesangial cell and podocyte injuries in diabetic nephropathy. Nephrology 23, 32–37 (2018).

12. Kamiya, N. et al. Wnt inhibitors Dkk1 and sost are downstream targets of BMP signaling through the type IA receptor (BMPRIA) in osteoblasts. J. Bone Miner. Res. 25, 200–210 (2010).

13. Zhou, L. et al. Mutual antagonism of Wilms’ tumor 1 and $β$-catenin dictates podocyte health and disease. J. Am. Soc. Nephrol. 26, 677–691 (2015).

14. Nishinakamura, R. & Sakaguchi, M. BMP signaling and its modifiers in kidney development. Pediatr. Nephrol. 29, 681–686 (2014).

15. Lindström, N. O. et al. Conserved and Divergent Molecular and Anatomic Features of Human and Mouse Nephron Patterning. J. Am. Soc. Nephrol. 29, 825–840 (2018).

16. Nakai, S. et al. Crucial roles of Brn1 in distal tubule formation and function in mouse kidney. Development 130, 4751–4759 (2003).

17. Massa, F. et al. Hepatocyte nuclear factor 1β controls nephron tubular development. Dev. 140, 886–896 (2013).

18. Naylor, R. W., Przepiorski, A., Ren, Q., Yu, J. & Davidson, A. J. HNF1$β$ is essential for nephron segmentation during nephrogenesis. J. Am. Soc. Nephrol. 24, 77–87 (2013).

19. Deacon, P., Concodora, C. W., Chung, E. & Park, J. S. Β-Catenin Regulates the Formation of Multiple Nephron Segments in the Mouse Kidney. Sci. Rep. 9, 1–13 (2019).

20. O’Brien Lori L., M. A.P. Induction and patterning of the metanephric nephron. Semin cell dev biol 0, 31–38 (2014).

21. Chung, E., Deacon, P. & Park, J. S. Notch is required for the formation of all nephron segments and primes nephron progenitors for differentiation. Dev. 144, 4530–4539 (2017).

22. Cheng, H. T. et al. Erratum: Notch2, but not Notch1, is required for proximal fate acquisition in the mammalian nephron (Development vol. 134 (801-811)). Development 134, 4506 (2007).

23. Park, J. S., Valerius, M. T. & McMahon, A. P. Wnt/β-catenin signaling regulates nephron induction during mouse kidney development. Development 134, 2533–2539 (2007).

24. Bugacov, H. et al. Dose-dependent responses to canonical Wnt transcriptional complexes in the regulation of mammalian nephron progenitors. Development 151, (2024).

25. Der, B., Bugacov, H., Briantseva, B.-M. & McMahon, A. P. Cadherin Adhesion Complexes Direct Cell Aggregation in the Epithelial Transition of Wnt-Induced Nephron Progenitor Cells. bioRxiv Prepr. Serv. Biol. (2023) doi:10.1101/2023.08.27.555021.

26. Carroll, T. J., Park, J. S., Hayashi, S., Majumdar, A. & McMahon, A. P. Wnt9b plays a central role in the regulation of mesenchymal to epithelial transitions underlying organogenesis of the mammalian urogenital system. Dev. Cell 9, 283–292 (2005).

27. Karner, C. M. et al. Canonical Wnt9b signaling balances progenitor cell expansion and differentiation during kidney development. Development 138, 1247–57 (2011).

28. Lindström, N. O. et al. Integrated β-catenin, BMP, PTEN, and Notch signalling patterns the nephron. Elife 3, e04000 (2015).

29. Takasato, M. & Little, M. H. A strategy for generating kidney organoids: Recapitulating the development in human pluripotent stem cells. Dev. Biol. 420, 210–220 (2016).

30. Ueda, H. et al. Bmp in podocytes is essential for normal glomerular capillary formation. J. Am. Soc. Nephrol. 19, 685–94 (2008).

31. Ng-Blichfeldt, J.-P., Stewart, B. J., Clatworthy, M. R., Williams, J. M. & Röper, K. Identification of a core transcriptional program driving the human renal mesenchymal-to-epithelial transition. Dev. Cell 59, 595–612.e8 (2024).

32. Yu, P. B. et al. BMP type I receptor inhibition reduces heterotopic ossification. Nat. Med. 14, 1363–1369 (2008).

33. Schnell, J., et al. Stepwise developmental mimicry generates proximal-biased kidney organoids. bioRxiv (2024).

34. Liu, Z. et al. The extracellular domain of notch2 increases its cell-surface abundance and ligand responsiveness during kidney development. Dev. Cell 25, 585–598 (2013).

35. Naiman, N. et al. Repression of Interstitial Identity in Nephron Progenitor Cells by Pax2 Establishes the Nephron-Interstitium Boundary during Kidney Development. Dev. Cell 41, 349–365.e3 (2017).

36. Wilson, S. B. & Little, M. H. The origin and role of the renal stroma. Development 148, (2021).

37. Inoue, S., Inoue, M., Fujimura, S. & Nishinakamura, R. A mouse line expressing Sall1-driven inducible Cre recombinase in the kidney mesenchyme. Genesis 48, 207–12 (2010).

38. Ying, Q. L., Nichols, J., Chambers, I. & Smith, A. BMP induction of Id proteins suppresses differentiation and sustains embryonic stem cell self-renewal in collaboration with STAT3. Cell 115, 281–92 (2003).

39. Kim, A. D. et al. Cellular Recruitment by Podocyte-Derived Pro-migratory Factors in Assembly of the Human Renal Filter. iScience 20, 402–414 (2019).

40. Böhm, J., Sustmann, C., Wilhelm, C. & Kohlhase, J. SALL4 is directly activated by TCF/LEF in the canonical Wnt signaling pathway. Biochem. Biophys. Res. Commun. 348, 898–907 (2006).

41. Ito, T. et al. Potential role of the OVOL1-OVOL2 axis and c-Myc in the progression of cutaneous squamous cell carcinoma. Mod. Pathol. 30, 919–927 (2017).

42. Qiu, D. et al. Klf2 and Tfcp2l1, Two Wnt/β-Catenin Targets, Act Synergistically to Induce and Maintain Naive Pluripotency. Stem cell reports 5, 314–22 (2015).

43. Hofmann, M. et al. WNT signaling, in synergy with T/TBX6, controls Notch signaling by regulating Dll1 expression in the presomitic mesoderm of mouse embryos. Genes Dev. 18, 2712–2717 (2004).

44. Miao, Z. et al. Single cell regulatory landscape of the mouse kidney highlights cellular differentiation programs and disease targets. Nat. Commun. 12, (2021).

45. Yu, P. B. et al. Dorsomorphin inhibits BMP signals required for embryogenesis and iron metabolism. Nat. Chem. Biol. 4, 33–41 (2008).

46. Hao, J. et al. Dorsomorphin, a selective small molecule inhibitor of BMP signaling, promotes cardiomyogenesis in embryonic stem cells. PLoS One 3, e2904 (2008).

47. Emerling, B. M. et al. Depletion of a putatively druggable class of phosphatidylinositol kinases inhibits growth of p53-null tumors. Cell 155, 844–57 (2013).

48. Vanslambrouck, J. M. et al. A toolbox to characterize human induced pluripotent stem cell-derived kidney cell types and organoids. J. Am. Soc. Nephrol. 30, 1811–1823 (2019).

49. Tran, T. et al. In Vivo Developmental Trajectories of Human Podocyte Inform In Vitro Differentiation of Pluripotent Stem Cell-Derived Podocytes. Dev. Cell 50, 102–116.e6 (2019).

50. Lake, B. B. et al. An atlas of healthy and injured cell states and niches in the human kidney. Nature 619, 585–594 (2023).

51. Grieshammer, U. et al. FGF8 is required for cell survival at distinct stages of nephrogenesis and for regulation of gene expression in nascent nephrons. Development 132, 3847–3857 (2005).

52. Fausto, C. C. et al. Defining and Controlling Axial Nephron Patterning in Human Kidney Organoids with Synthetic Wnt-Secreting Organizers. (2024) doi:10.1101/2024.11.30.626171.

53. Yamashita, T., Konishi, M., Miyake, A., Inui, K. & Itoh, N. Fibroblast growth factor (FGF)-23 inhibits renal phosphate reabsorption by activation of the mitogen-activated protein kinase pathway. J. Biol. Chem. 277, 28265–70 (2002).

54. Sato, A. et al. FGF8 signaling is chemotactic for cardiac neural crest cells. Dev. Biol. 354, 18–30 (2011).

55. Heliot, C. et al. HNF1B controls proximal-intermediate nephron segment identity in vertebrates by regulating Notch signalling components and Irx1/2. Development 140, 873–885 (2013).

56. Massa, F. M. et al. MAFB drives differentiation by permitting WT1 binding to podocyte specific promoters. (2023) doi:10.7554/eLife.93138.1.

57. Dong, L. et al. Integration of Cistromic and Transcriptomic Analyses Identifies Nphs2, Mafb, and Magi2 as Wilms’ Tumor 1 Target Genes in Podocyte Differentiation and Maintenance. J. Am. Soc. Nephrol. 26, 2118–28 (2015).

58. Kann, M. et al. Genome-Wide Analysis of Wilms’ Tumor 1-Controlled Gene Expression in Podocytes Reveals Key Regulatory Mechanisms. J. Am. Soc. Nephrol. 26, 2097–104 (2015).

59. Harshuk-Shabso, S., Dressler, H., Niehrs, C., Aamar, E. & Enshell-Seijffers, D. Fgf and Wnt signaling interaction in the mesenchymal niche regulates the murine hair cycle clock. Nat. Commun. 11, 5114 (2020).

60. Kjolby, R. A. S., Truchado-Garcia, M., Iruvanti, S. & Harland, R. M. Integration of Wnt and FGF signaling in the Xenopus gastrula at TCF and Ets binding sites shows the importance of short-range repression by TCF in patterning the marginal zone. Development 146, (2019).

61. Cheng, H.-T. et al. Notch2, but not Notch1, is required for proximal fate acquisition in the mammalian nephron. Development 134, 801–11 (2007).

62. Boyle, S. C., Kim, M., Valerius, M. T., Mcmahon, A. P. & Kopan, R. Notch pathway activation can replace the requirement for Wnt4 and wnt9b in mesenchymal-to-epithelial transition of nephron stem cells. Development 138, 4245–4254 (2011).

63. Yaman, Y. I. & Ramanathan, S. Controlling human organoid symmetry breaking reveals signaling gradients drive segmentation clock waves. Cell 186, 513–527.e19 (2023).

64. Anand, G. M. et al. Controlling organoid symmetry breaking uncovers an excitable system underlying human axial elongation. Cell 186, 497–512.e23 (2023).

65. Atamian, A. et al. Human cerebellar organoids with functional Purkinje cells. Cell Stem Cell 31, 39–51.e6 (2024).

66. Vanslambrouck, J. M. et al. Enhanced metanephric specification to functional proximal tubule enables toxicity screening and infectious disease modelling in kidney organoids. Nat. Commun. 13, 5943 (2022).

67. Schneider, J., Arraf, A. A., Grinstein, M., Yelin, R. & Schultheiss, T. M. Wnt signaling orients the proximal-distal axis of chick kidney nephrons. Dev. 142, 2686–2695 (2015).

68. Low, J. H. et al. Generation of Human PSC-Derived Kidney Organoids with Patterned Nephron Segments and a De Novo Vascular Network. Cell Stem Cell 25, 373–387.e9 (2019).

69. Shi, M. et al. Integrating collecting systems in kidney organoids through fusion of distal nephron to ureteric bud. bioRxiv Prepr. Serv. Biol. (2024) doi:10.1101/2024.09.19.613645.

70. Mugford, J. W., Yu, J., Kobayashi, A. & McMahon, A. P. High-resolution gene expression analysis of the developing mouse kidney defines novel cellular compartments within the nephron progenitor population. Dev. Biol. 333, 312–23 (2009).

71. Park, J. S. et al. Six2 and Wnt Regulate Self-Renewal and Commitment of Nephron Progenitors through Shared Gene Regulatory Networks. Dev. Cell 23, 637–651 (2012).

72. Schnell, J., Achieng, M. A. & Lindström, N. O. Principles of human and mouse nephron development. Nat. Rev. Nephrol. 0123456789, (2022).

73. Duvall, K. et al. Revisiting the role of Notch in nephron segmentation confirms a role for proximal fate selection during mouse and human nephrogenesis. Development 149, (2022).

74. Huang, B. et al. Long-term expandable mouse and human-induced nephron progenitor cells enable kidney organoid maturation and modeling of plasticity and disease. Cell Stem Cell 31, 921–939.e17 (2024).

75. Lindström, N. O. et al. Conserved and Divergent Features of Mesenchymal Progenitor Cell Types within the Cortical Nephrogenic Niche of the Human and Mouse Kidney. J. Am. Soc. Nephrol. 29, 806–824 (2018).

76. Lindström, N. O. et al. Conserved and divergent features of human and mouse kidney organogenesis. J. Am. Soc. Nephrol. 29, 785–805 (2018).

77. O’Rahilly, R., Müller, F., Hutchins, G. M. & Moore, G. W. Computer ranking of the sequence of appearance of 73 features of the brain and related structures in staged human embryos during the sixth week of development. Am. J. Anat. 180, 69–86 (1987).

78. O’Rahilly, R. & Müller, F. Developmental stages in human embryos: revised and new measurements. Cells. Tissues. Organs 192, 73–84 (2010).

79. Strachan, T., Lindsay, S. & Wilson, D. I. Molecular Genetics of Early Human Development. (BIOS Scientific, 1997).

80. Takasato, M., Er, P. X., Chiu, H. S. & Little, M. H. Generation of kidney organoids from human pluripotent stem cells. Nat. Protoc. 11, 1681–92 (2016).

81. Wörsdörfer, P., Rockel, A., Alt, Y., Kern, A. & Ergün, S. Generation of Vascularized Neural Organoids by Co-culturing with Mesodermal Progenitor Cells. STAR Protoc. 1, 100041 (2020).

82. Dobin, A. et al. STAR: ultrafast universal RNA-seq aligner. Bioinformatics 29, 15–21 (2013).

83. Liao, Y., Smyth, G. K. & Shi, W. The R package Rsubread is easier, faster, cheaper and better for alignment and quantification of RNA sequencing reads. Nucleic Acids Res. 47, e47 (2019).

84. Love, M. I., Huber, W. & Anders, S. Moderated estimation of fold change and dispersion for RNA-seq data with DESeq2. Genome Biol. 15, 550 (2014).

85. Stuart, T. et al. Comprehensive Integration of Single-Cell Data. Cell 177, 1888–1902.e21 (2019).

86. Nestorowa, S. et al. A single-cell resolution map of mouse hematopoietic stem and progenitor cell differentiation. Blood 128, e20–31 (2016).

87. Hafemeister, C. & Satija, R. Normalization and variance stabilization of single-cell RNA-seq data using regularized negative binomial regression. Genome Biol. 20, 296 (2019).

88. Ng, A. H. M. et al. A comprehensive library of human transcription factors for cell fate engineering. Nat. Biotechnol. 39, 510–519 (2021).

89. Zhou, Q. et al. A mouse tissue transcription factor atlas. Nat. Commun. 8, 15089 (2017).

90. Cunningham, F. et al. Ensembl 2022. Nucleic Acids Res. 50, D988–D995 (2022).

91. Hao, Y. et al. Integrated analysis of multimodal single-cell data. Cell 184, 3573–3587.e29 (2021).

